# Suite3D: Volumetric cell detection for two-photon microscopy

**DOI:** 10.1101/2025.03.26.645628

**Authors:** Ali Haydaroğlu, Tinya Chang, Andrew Landau, Michael Krumin, Suyash Agarwal, Sam Dodgson, Liad J. Baruchin, Maria Cozan, Jingkun Guo, David Meyer, Charu Bai Reddy, Jian Zhong, Na Ji, Sylvia Schröder, Kenneth D Harris, Alipasha Vaziri, Matteo Carandini

**Affiliations:** University College London, London, UK; Beth Israel Deaconess Medical Center, Harvard Medical School, Boston, MA, USA; University of Sussex, Brighton, UK; Rockefeller University, New York, NY, USA; University of California Berkeley, CA, USA

## Abstract

In two-photon imaging of neuronal activity it is increasingly common to acquire 3-dimensional volumes. However, these volumes are typically processed plane by plane, leading to uncorrected axial movement, duplicated cells across planes, reduced signal-to-noise ratio per cell, and missed cells. To overcome these limitations, we introduce Suite3D, a volumetric cell detection pipeline. Suite3D corrects for rigid and non-rigid 3D brain motion, both lateral and axial. It detects neurons using 3D correlation, improving detectability. Finally, it performs 3D segmentation, identifying cells across imaging planes. We validated Suite3D with data from conventional multi-plane microscopes and advanced volumetric microscopes, at multiple resolutions and in multiple brain regions, and with ground truth anatomical labelling. Suite3D successfully detected cells appearing on multiple imaging planes, improving cell detectability and signal quality, avoiding duplications, and running faster than a prior volumetric pipeline. Suite3D offers a powerful solution for analyzing volumetric two-photon data.

---

The wide adoption of 2-photon microscopes has made it common to image 3D volumes of brain tissue, greatly increasing the number of neurons that are sampled in the imaged region. With conventional 2-photon microscopes, volumetric imaging is typically achieved by scanning each plane in sequence^1^. This approach has yielded recordings from increasing numbers of neurons, from over 1,000 neurons in a 0.03 mm^3^ volume^2^ to over 10,000 neurons in a 0.3 mm^3^ volume^3^. These numbers can be further increased with imaging systems that use temporal multiplexing approaches to obtain volumetric imaging. One such approach is Free-space Angular-Chirp-Enhanced Delay (FACED) imaging, which turns a light pulse into an ultrafast linear sequence of pulses, enabling passive line scanning^4–6^. Another approach is Light Beads Microscopy (LBM), which turns a light pulse into a column of light beads at multiple depths, thus acquiring an entire volume while scanning only one plane^7^. This approach has the potential of recording close to 1,000,000 neurons in a 15 mm^3^ volume^7,8^. Beyond 2-photon imaging, methods such as light-sheet microscopy^9^, light-field microscopy^10^, and SCAPE^11,12^ are capable of fast, high-resolution volumetric imaging of *in vivo* neural activity.

These imaging methods produce vast volumetric datasets, which are typically processed one plane at a time or segmented manually^2,5^. Processing 2-photon data requires a pipeline that performs a sequence of operations from motion correction to cell segmentation. Established pipelines such as Suite2p^13,14^, 2D CaImAn^15–17^, FIOLA^18^, and EXTRACT^19^ perform this sequence of operations independently on individual planes. They are typically used even if the data are volumetric^3,8^, sometimes with post-hoc corrections to merge cells that appear in multiple imaging planes^7,20^.

By contrast, efforts to extend the processing to volumes, such as volumetric CaImAn^15,21^, have been met with limited adoption perhaps due to their extensive computational demands. Volumetric segmentation is commonly performed in anatomical datasets with tools such as CellPose^22^ and CellSeg3D^23^, but these tools ignore the time course of the activity, missing out on a powerful aid to detection and segmentation.

When applied to volumetric data, however, plane-by-plane methods can lead to multiple problems, including cell duplications, signal losses, and missed neurons. First, if a cell has significant signal on two or more planes, it will be split and detected independently in each of those planes, causing duplications. Second, when a cell is split across planes, the region detected on each plane will have fewer pixels and therefore lower signal than the full cell. Third, a cell may be below the detection threshold on one (or all) individual planes, and only detectable when planes are analyzed together. Finally, brain movement perpendicular to the imaging plane cannot be corrected with plane-by-plane methods, and it can have strong effects that can be mistaken for functional signals, especially when imaging small structures^24^.

Here we show that these concerns apply to many 2-photon datasets, including those obtained with conventional 2-photon microscopes, and we resolve them with Suite3D, a fully volumetric motion correction and cell detection pipeline. Suite3D extends to 3 dimensions the 2D pipeline of Suite2p, with novel 3D algorithms, computational improvements, and visualization tools. It is faster than plane-by-plane methods, and runs dramatically faster than volumetric CaImAn. Its algorithms are robust to variations in signal levels and are tunable, with intuitive parameter selection and with visualization interfaces for semi-automated curation.

We validated Suite3D with data acquired with conventional 2-photon microscopes and with advanced volumetric microscopes. Suite3D successfully detected cells appearing on multiple imaging planes, improving their detectability, signal quality, and identification. By contrast, plane-by-plane segmentation led to duplication of cells and to reduced SNR. Suite3D is thus a promising tool for analyzing 2-photon volumes acquired with a range of microscopy approaches.

## Results

We begin by illustrating how in a typical 2-photon imaging volume neurons often straddle multiple imaging planes. We then show how this issue can be addressed and indeed exploited through volumetric processing. We describe Suite3D, compare it to other popular approaches, and apply it to a variety of datasets.

### Neurons straddle imaging planes

To illustrate how common it is for neurons to straddle imaging planes, we acquired a typical multi-plane dataset with a conventional 2-photon microscope (Figure 1a) and analyzed the results both with the common plane-by-plane analysis approach (Figure 1 b) and with our volumetric approach (Figure 1c). We imaged Layer 5 (L5) neurons in the mouse visual cortex and acquired multiple planes separated by 20 µm, which is in the range commonly used in the field. In this mouse, L5 excitatory neurons expressed the green functional indicator GCaMP6s (Figure 1d), and a sparser subset of neurons also expressed the red anatomical marker tdTomato (Figure 1e). We can thus be sure that when red pixels appear at similar positions across planes, they correspond to the same neuron.

**Figure 1.**
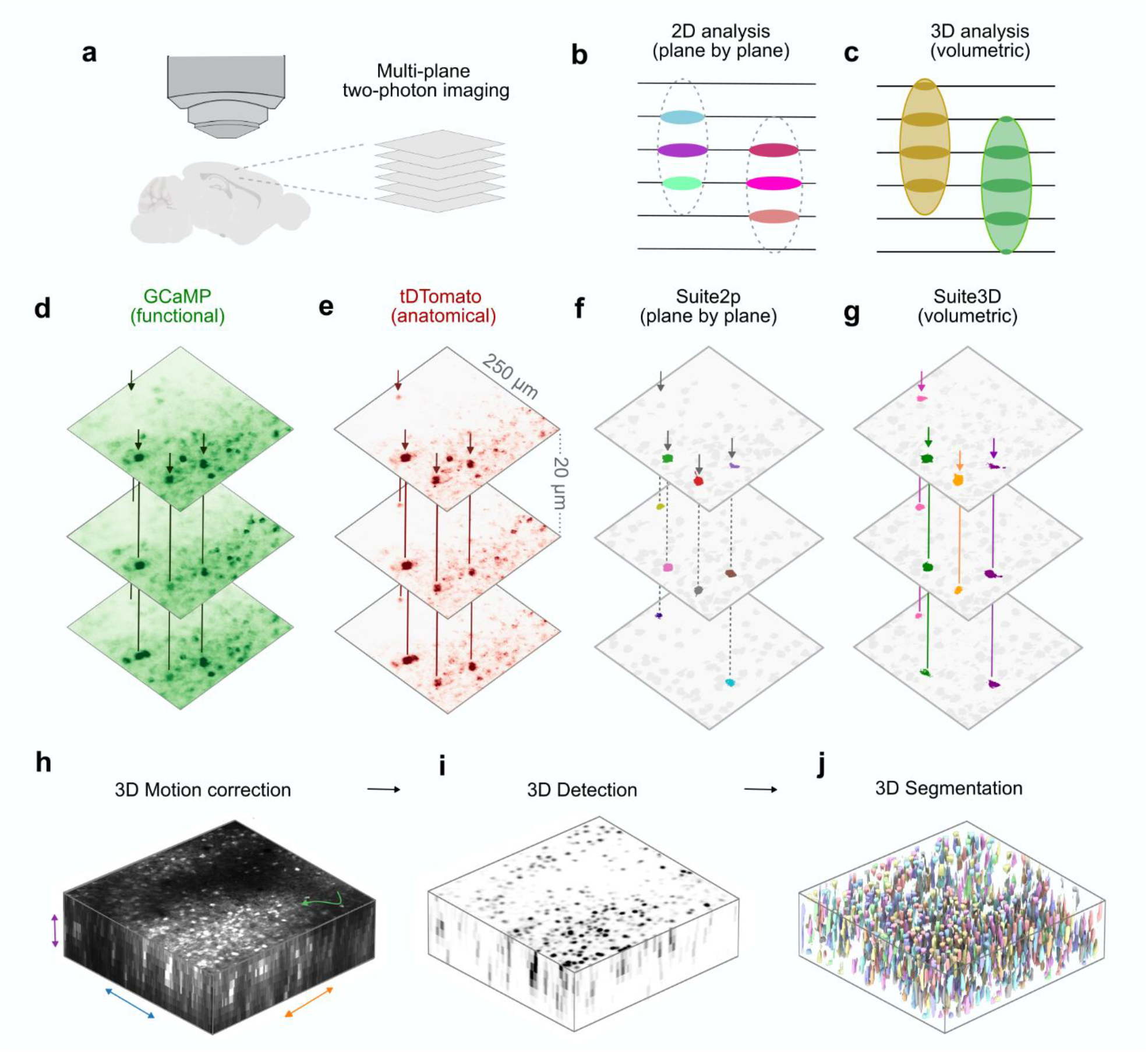
Neurons straddle imaging planes, requiring a 3D analysis pipeline. **a.** Multi-plane imaging methods record volumetric data. **b**. Cartoon of a z-section from a volumetric dataset analyzed with plane-by-plane analysis methods. Two cells that straddle planes (*ellipses*) are given different identities on different planes (*colors*) and are occasionally missed. **c**. Same as (b), analyzed volumetrically. The two cells are identified and all their pixels are counted. **d,e**. Mean fluorescence images from three planes of an example multi-plane recording in Layer 5 of mouse visual cortex with excitatory cells expressing GCaMP, and a sparse subset expressing tdTomato. Arrows indicate 4 example cells that straddle planes. **f**. Plane-by-plane analysis on the example dataset identified some of the pixels of the 4 example cells, assigning multiple identities to them (colors) and missing some of their pixels in some of the planes. **g**. Analysis with Suite3D identified all of the example cells in all planes. **h**. The first stage in Suite3D is 3D motion correction, which corrects for rigid and non-rigid motion in all three axes. **i**. The second stage is 3D detection, which produces a correlation volume. **j**, The third stage is 3D segmentation, which assigns voxels to 3D regions of interest (ROIs).

The results revealed that neurons commonly appear in two or more planes (Figure 1d,e). This observation should perhaps not be surprising, because neurons in the mouse cortex have somas that easily exceed ~20 µm in diameter^25,26^ and pyramidal neurons may extend more in the axial dimension due to their apical dendrites. Moreover, the 2-photon point spread function is more extended in the axial than lateral dimension, and is typically on the order of 10 µm (Ref. ^27^).Together, these factors mean that fluorescence from single neurons can cross multiple imaging planes even if these planes are ≥30 µm apart.

And yet, plane-by-plane analyses of volumetric datasets typically disregard the possibility that neurons could straddle planes, leading to duplications and to inefficient assignment of pixels to neurons. For instance, analyzing the example dataset plane-by-plane with Suite2p^13,14^ leads to a myriad duplications and to missed assignments of pixels to neurons (Figure 1f).

### A 3D analysis pipeline

To overcome these limitations, we developed Suite3D, a volumetric cell detection pipeline that operates simultaneously across planes. Suite3D resolves the limitations of plane-by-plane analyses. For instance, in the example dataset it correctly identified cells across planes, and assigned to neurons pixels that would otherwise have gone unassigned (Figure 1g).

Suite3D comprises three core processing stages. The first stage is 3D motion correction, which corrects for lateral and axial brain movements (Figure 1h). The second stage is 3D detection, through spatiotemporal filtering, normalization, and thresholding (Figure 1i). The third stage is 3D segmentation, where voxels are assigned to distinct regions of interest (ROIs, Figure 1j). Below we describe these steps in order.

### 3D motion correction

The first stage in the Suite3D pipeline corrects for brain motion, computing this correction volumetrically rather than plane-by-plane as in standard pipelines. By operating volumetrically, Suite3D can estimate lateral shifts that are coherent across planes. Moreover, it can estimate axial shifts, which would be undetectable in individual planes. This correction is applied in two steps: first, a rigid correction applied to the entire volume, next, a non-rigid correction applied to smaller 3D blocks to correct for local movement that may not be shared by the rest of the volume.

The volumetric motion correction is based on phase correlation. For the first step, rigid motion correction, we took the basic idea from Suite2p^13,14^, extended it to 3D, and GPU-accelerated it to make it tractable. At each timestep, Suite3D computes the 3D Fourier transform of the acquired volume and its 3D phase correlation with a reference volume. From the 3D phase correlation, it then computes x-, yand z-shifts, and applies them to the acquired volume to correct motion. For the second step, non-rigid correction, Suite3D divides the acquired volume in smaller 3D blocks and applies the same GPU-accelerated procedure to each block^28^.

This 3D motion correction is markedly superior to plane-by-plane methods. To compare the two methods, we analyzed the same volume with 2D and 3D motion correction. The volume was acquired with a Light Beads Microscope^7^ (LBM), which images multiple planes (placed 20 µm apart) near-simultaneously (8 ns delay). The plane-by-plane motion correction (Figure 2a) often estimated different shifts even for nearby planes (Figure 2b). This disparity across planes is unlikely to reflect genuine shearing of brain tissue, as the planes are spaced only 20 µm apart and acquired near-simultaneously. Instead, it is due to the low signal-to-noise ratio in the phase correlation maps, which makes it difficult to estimate their peaks in the presence of noise (Figure 2c). By contrast, the 3D motion correction implemented by Suite3D (Figure 2e) produced more robust motion estimates, and it did so in three dimensions instead of two (Figure 2f), with much sharper phase correlation maps (Figure 2g). The sharpness of the phase correlation maps is quantified by the signal-to-noise ratio metric^13^, which is much higher for 3D than 2D motion estimation (Figure 2i). This superiority is not due to the 3D approach being nonrigid and the 2D approach being rigid: Similar results were obtained when the 3D approach was limited to being rigid (ED Figure 1).

**Figure 2.**
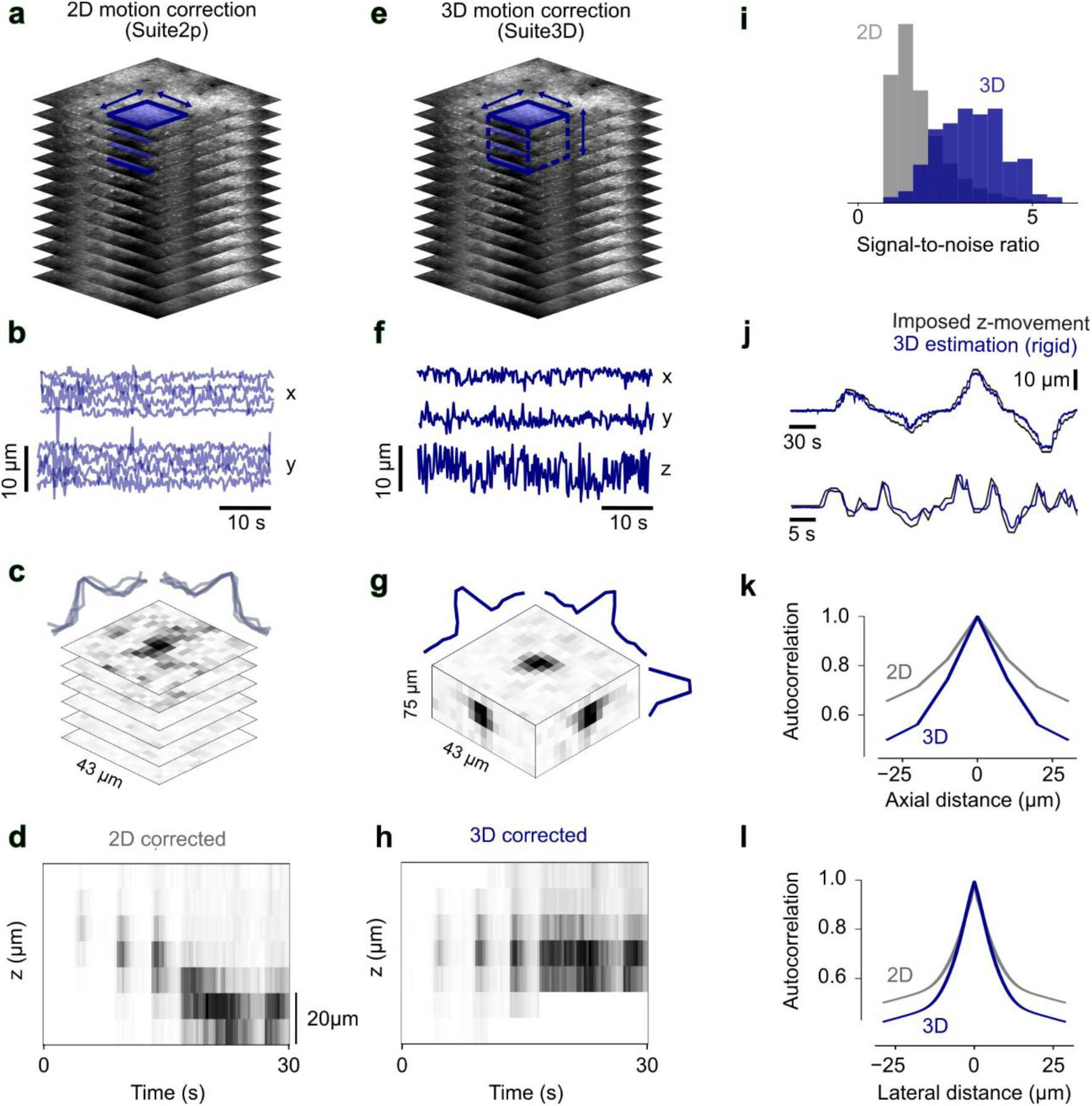
3D motion correction. Nonrigid 3D motion correction improves planar shift estimates over 2D motion correction, and corrects for axial motion. **a.** Reference image for an example LBM recording, highlighting the planar patches (*blue*) that are shifted independently in 2D motion correction, correcting only for within-plane motion. **b**. Standard 2D motion correction estimates x and y shifts independently for each plane, leading to different estimated shifts across planes. **c**. Phase correlation for an example frame for the patches in b. The overlaid curves show the center slice for each patch. **d**. Activity over time in a column of pixels centered on a neuron during induced z-shift, which cannot be corrected by 2D motion correction. **e-h**. Same as a-d, for 3D motion correction, which estimates three-dimensional motion for blocks of the volume (*blue* cube in e), resulting in a single estimate of x, y and z shifts for the block (f), with tight phase correlations across the block (g) and successful motion correction in the z plane (h). **i**, Distribution of the signal-to-noise ratio in the phase correlation maps (c,g) across patches (2D) and blocks (3D). **j**. In a recording with imposed z-movement (same as panels d and h), 3D motion correction recovers the ground truth movement. **k**. Spatial autocorrelation in the axial direction of the motion-corrected movie. **l**. Same, in the lateral direction. For equivalent results with rigid volumetric motion correction see ED Figure 1.

Moreover, Suite3D accurately estimates and corrects for axial motion. To test Suite3D’s ability to estimate axial motion, we used a conventional multi-plane 2p microscope to image a 7plane volume spanning 120 µm in depth, and we introduced intentional axial motion by moving the objective by as much as 40 µm while recording (Figure 2j). Suite3D passed this test with flying colors, accurately reconstructing and correcting this ground-truth axial motion (Figure 2h,j).

By contrast, 2D detection pipelines such as Suite2p cannot detect axial motion, but rather suffer from its consequences. Indeed, axial motion can lead to cells shifting in and out of plane (Figure 2d). This shifting will lower the signal/noise ratio, and if it is systematic (e.g. if it correlates with behavior), it may even confound the functional characterization of cells. Suite3D, instead, prevents this confound by successfully correcting the cell drift across planes (Figure 2h).

These abilities of Suite3D’s motion correction stage allow the subsequent stages to operate on sharper images. Motion blur reduces the sharpness of images^24^, broadening their spatial autocorrelation. This autocorrelation was substantially sharper after correcting for motion volumetrically with Suite3D than after correcting plane-by-plane with Suite2p, both laterally (Figure 2k) and axially (Figure 2l).

To perform these operations efficiently, Suite3D exploits “pipelining” and GPU acceleration. Volumetric microscopy produces large data sets – for instance, the LBM produces >700 GB/hour. Therefore, it is often unfeasible to store raw data locally on the processing computer, and data loading becomes a major bottleneck. Suite3D substantially reduces this problem by pipelining: it uses a separate thread to load the next batch from disk while the current batch is being processed. In addition, it accelerates the Fourier transform and phase correlation by performing them on a GPU with custom CUDA kernels.

### 3D detection

The next stage in the Suite3D pipeline is to detect regions of interest (ROIs) through spatio-temporal processing steps that convert the (4D) motion-corrected movie – a sequence of volumes – into a 3D “correlation volume”, which will be used in the final step: 3D segmentation (Figure 3a-d, Suppl. Video 1).

**Figure 3.**
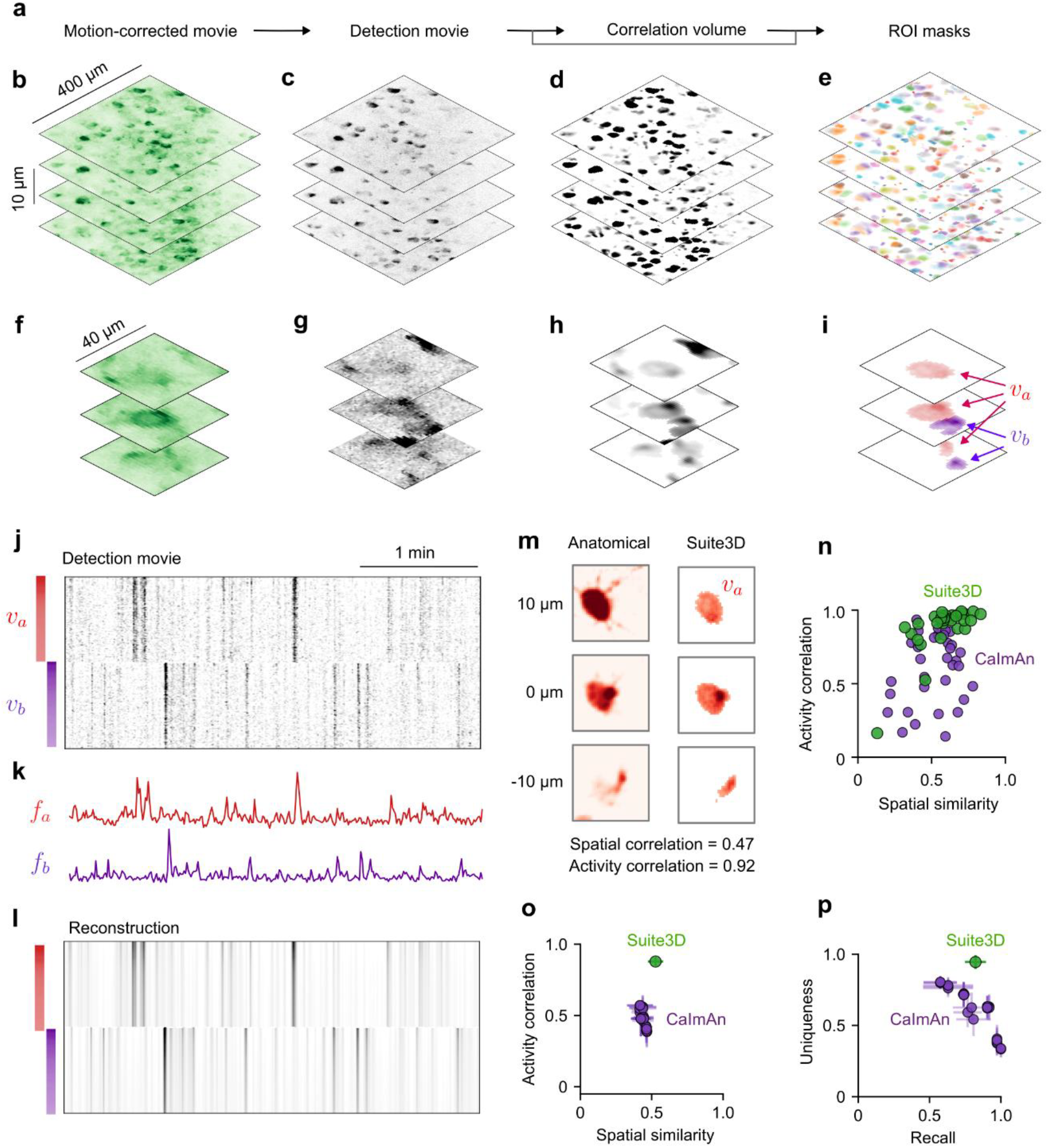
3D detection and segmentation. **a**, The main steps in detection and segmentation. **b**, Mean image of the motion-corrected movie for an example multi-photon 2p dataset. **c.** Standard deviation image of the detection movie for the same dataset. **d**. Correlation volume for the same dataset. **e**. ROI masks from Suite3D for the same dataset. **f-i**, Zoomed-in view of the same dataset, in a volume where Suite3D found two ROIs, whose volumes are denoted v_a_ and v_b_ (panel i). **j**. Activity (from the detection movie) in the top voxels of the two ROIs, with spatial weights plotted on the left (*color bars*). **k**. Time courses of the detection movie in the two ROIs. **l**. Reconstruction of the voxel activity from an outer product of the spatial weights of each voxel and the time course of activity in each ROI. **m**. Anatomical validation of the neuron denoted as v_a_ (*red* in panel i), showing the ground truth (tdTomato) label of the neuron in each plane (*left*) and the corresponding Suite3D masks (*right*). The spatial similarity between the two is 0.47, and the temporal correlation between the corresponding traces is 0.92. **n**, Spatial similarity and activity correlation with the ground truth ROIs for ROIs obtained with Suite3D (*green*) and with CaImAn (*purple*). **o**. Same, averaged over all cells in three mice. Each purple point corresponds to one of 36 parameter configurations of CaImAn. Error bars indicate s.e. across mice (n=3). **p**. Fraction of detected cells (recall) and uniqueness of detected cells based on ground truth labels, for Suite3D (*green*) and CaImAn (*purple*).

The first step in this process is the computation of a (4D) “detection movie”. The motion-corrected movie (Figure 3b) is high-pass filtered in time (typically with a 0.1 Hz cutoff for slow calcium indicators) and each pixel is normalized by the standard deviation of its first temporal differences (Figure 3c). These steps exclude slow temporal drifts and normalize for differences in signal across the volume. The resulting movie is then high-pass filtered in space (typically at 70 μm) to subtract the neuropil signal.

The next step is the computation of a 3D “correlation volume” (Figure 3a). This volume highlights pixels that have high correlation to their neighbors at cell-like spatiotemporal scales (or at smaller scales if the targets are synapses, as we will see later). The procedure is similar to the 2D correlation map in Suite2p^14^, adapted to 3D and modified to be computationally tractable on large-scale datasets. The detection movie is first low-pass filtered to highlight cell-sized structures (or smaller structures, as needed). This filtered movie is then reduced to a single 3D volume. To achieve this reduction, one could take the mean or root-mean-square of each pixel over time; however, if a cell were to only activate a few times in a long recording, it would be hard to detect. Instead, we threshold each pixel in the movie before taking the root-sum-square and thus obtain a “correlation volume” (Figure 3d).

The correlation volume is particularly useful in cases where neurons have dim fluorescence or sparse activity. For example, consider a neuron whose GCaMP fluorescence is mostly concentrated on two planes (Figure 3f). The detection movie separates that neuron from the background (Figure 3g) and the correlation volume shows evidence of correlated activity on an additional plane (Figure 3h). As we will see shortly, this inference is confirmed by the anatomical ground-truth for this neuron.

The 3D correlation volume improves cell detectability compared to 2D methods. Volumetric data contain axial correlations^29^ that would be lost if one computed 2D correlation maps for each plane^13,14^ and stitched them together into a volume (ED Figure 2). Indeed, when applied to the same dataset, the 2D correlation maps had substantially more background noise than the 3D correlation volume (ED Figure 3). Moreover, cells with small footprints on a given plane would not be detectable in a 2D correlation map. Rather than contributing to the signal of the imaged neuron, those pixels would contribute to the background noise.

To make these steps feasible on vast datasets (several TB), Suite3D uses parallelization and memory mapping. It also includes an intuitive parameter-sweeping interface that allows the user to systematically tune parameters and improve the correlation volume.

### 3D segmentation

The final stage in the Suite3D pipeline is 3D segmentation (Figure 3e). Similar to the 2D algorithm in Suite2p^14^, Suite3D performs segmentation as an iterative process. First, peaks in the correlation volume are used to seed the ROIs. Then, each ROI is grown based on its activity in the detection movie. All the voxels that neighbor an ROI are evaluated for their similarity in activity with the ROI activity, using a metric called voxel-SNR (Methods). The voxels where similarity is above a threshold are included in the ROI. The process is then repeated for each ROI until the ROI stops growing. This method approximates a low-rank decomposition of the detection movie with a locality constraint on the components (Figure 3 j-l).

3D segmentation detects cells that span multiple imaging planes (Figure 3e,i), which would be duplicated with 2D segmentation methods (ED Figure 3). Although post-hoc merging could potentially overcome this duplication, it is not trivial to merge 2D ROIs across planes. One implementation of post-hoc merging uses pairwise correlations of ROIs with nearby xy-coordinates on neighboring planes, and merges them if these correlations are above a threshold^7^. However, we show that the pairwise correlations of ROIs corresponding to the same cell can be very low, especially if an ROI has a small footprint on a given plane. The pairwise correlation of activity in ROIs belonging to the same cell falls well within the distribution of pairwise correlations of all ROIs within a recording, and a hard threshold will not sufficiently merge ROIs (ED Figure 3). Thus, using plane-by-plane segmentation methods in any multi-plane recording with dense spacing (<30 µm) can lead to over-counting of cells.

Instead of segmenting the full movie at once, Suite3D reduces the RAM requirement to typical levels (<64 GB) while maintaining processing speed by parallelizing across ROIs and movie patches, similar to methods implemented in CaImAn^15^. Extremely large movies (~ 1TB) which would otherwise be impossible to segment can be processed with Suite3D on a typical workstation. The volumetric implementation of the algorithm introduces several interpretable parameters and a tuning interface for optimizing the pipeline for challenging datasets (e.g. high-noise, low-resolution).

After segmentation, a time course of neural activity is obtained for each ROI by performing a weighted average, neuropil subtraction, and deconvolution. The mask of each ROI is used to compute a weighted average of the fluorescence of each ROI pixel from the motion-corrected movie as an average of its voxels weighted by the ROI mask. The neuropil fluorescence is then calculated in a hollow volumetric shell around each ROI (excluding nearby cells) and is subtracted from the ROI fluorescence, a volumetric extension of the Suite2p neuropil correction. The neuropil-subtracted fluorescence is then deconvolved with the OASIS algorithm to remove the effects of the indicator’s time course^30,31^.

### Anatomical and functional validation

In describing Suite3D’s stages of processing, we have highlighted some of its advantages over 2D methods in improving motion correction, cell detectability, and cell identification. We further quantified the advantages of Suite3D with ground-truth anatomical and functional validation.

As ground-truth validation we used recordings where a sparse subset of visual cortex neurons were labelled with a red anatomical marker (as in Figure 1). We compared the cells detected by Suite3D to the anatomical labels by correlating their spatial footprints and their corresponding time courses (Figure 3m-o). The Suite3D ROIs had a temporal correlation of 0.86 ± 0.03 with the ground truth cells (mean ± s.e., n=3 mice, Figure 3o). The spatial similarity averaged 0.49 ± 0.05, which is a high value considering that it compares the activity-weighted ROI masks with flat anatomical masks. Indeed, an example cell that was an excellent match had a spatial similarity of 0.47 (Figure 3m).

Suite3D also had good performance in identifying the ground-truth ROIs. It correctly detected 86 ± 7% (mean ± s.e., n=3 mice) of the red cells that were active during the recording (“recall”, Figure 3p). Since many green cells (expressing GCaMP) did not express the sparse red markers by design, we could not compute “precision”. Instead, we computed a “uniqueness” metric, which is 1 if a red cell overlaps with a single Suite3D ROI, and lower if it overlaps with more ROIs. Suite3D ROIs had a uniqueness score of 0.92 ± 0.05 (Figure 3p).

To put these results into perspective, we computed the same metrics for the volumetric version of the CaImAn algorithm. We ran that algorithm with 36 parameter configurations, and found Suite3D to outperform CaImAn across all configurations (Figure 3n-p).

We also compared the 3D ROIs detected by Suite3D with the 2D ROIs detected by Suite2p, and found that the latter tended to split cells across multiple planes, to miss the presence of cells on some planes, or to miss cells entirely (Figure 4a-c, ED Figure 3). To quantify the number of duplicates we computed the pairwise correlation of nearby ROIs and assigned a duplication score to each pair. The results confirm that Suite3D produces many fewer potential duplicates (Figure 4d).

**Figure 4.**
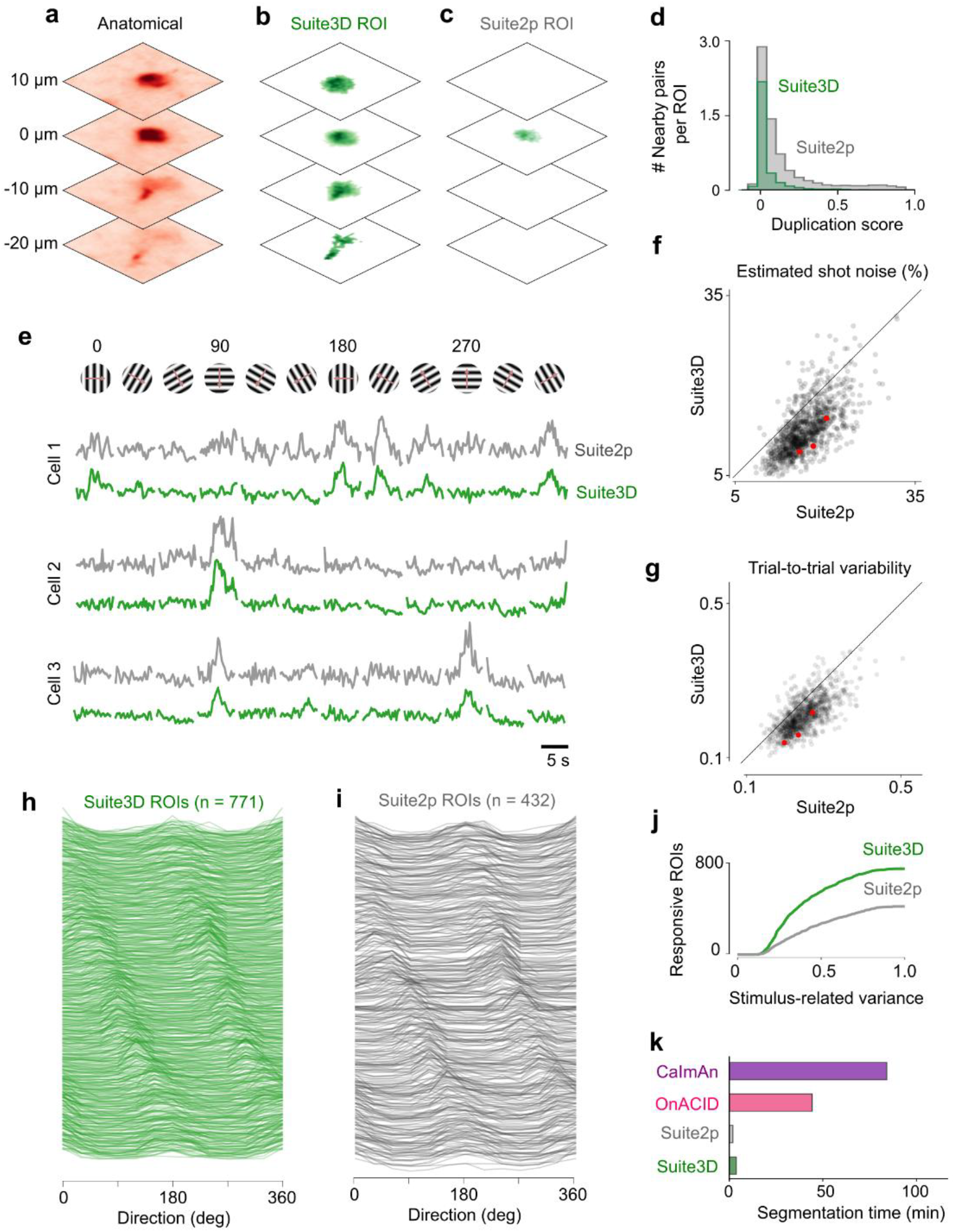
Anatomical and functional validation. **a**, Anatomical image of an example cell across four planes. **b.** Suite3D-detected mask spanning four planes for the cell in (a). **c**, Suite2p only finds an ROI in a single plane matching the cell in (a). **d**, Histogram of duplication score, computed only for pairs of cells nearby one another, normalized by the number of ROIs deected by Suite2p and Suite3D. **e**, dF/F traces of three example ROIs in visual cortex identified by both Suite2p and Suite3D in response to a single repeat of drifting gratings. **f**, For ROIs matched across the two pipelines, the estimated fraction of shot noise in the extracted fluorescence ^32^. Red dots indicate the cells shown in (e). **g**, Standard deviation of stimulus responses across 8 repeats for ROIs matched across the two pipelines. **h**, Direction tuning curves for all significantly stimulus-responsive ROIs found by Suite3D in an example session. **i**, Same as (h), for Suite2p ROIs. **j**, Cumulative histogram of stimulus-related variance for all stimulus-responsive ROIs found by each pipeline, computed on deconvolved traces. **k**, Runtime of Suite2p, Suite3D, CaImAn and OnACID to extract 1000 ROIs from the same recording, considering only the segmentation phase.

We next turned to functional measurements, and found that Suite3D improves the signal-to-noise ratio over Suite2p. We imaged the mouse visual cortex with a conventional 2p microscope while presenting drifting gratings of varying orientation and direction. We analyzed the resulting dataset with default parameters on both Suite3D and Suite2p and compared the raw fluorescence traces and deconvolved traces for ROIs detected by each pipeline. First, we compared cells that were identified in both pipelines by matching each Suite3D ROI with its closest matching planar Suite2p ROI. The dF/F traces extracted by Suite3D were smoother and appeared to have less shot noise (Figure 4e). This decrease in shot noise was confirmed by a quantitative estimator ^32^ (Methods) as being present in practically all detected cells (Figure 4f). To confirm this decrease corresponds to an increase in functional signal-to-noise ratio, we compared the standard deviation of dF/F across multiple repeats of the same stimulus for each cell, and found that the cells extracted by Suite3D were consistently more reliable (Figure 4g).

Remarkably, Suite3D also identified more stimulus-responsive cells (Figure 4h,i). The additional neurons found by Suite3D had a lower fraction of stimulus-related variance, suggesting that 2D methods miss neurons that are less responsive or noisier (Figure 4j). Nonetheless, the additional neurons had typical orientation tuning curves with two peaks separated by 180 degrees, and varied smoothly in their preferences (Figure 4h). Because of the salt-and-pepper organization of orientation tuning in mouse V1, such well-formed and diverse tuning curves are a positive assay of neuronal identification. Indeed, if neuronal signals were contaminated with neuropil signals, orientation tuning would appear weak and homogeneous. Moreover, if neurons that should be split are merged, the orientation tuning curves would likely have multiple peaks with odd spacings. The absence of these features is a further confirmation of Suite3D’s performance.

### Advantages over other 3D pipelines

We also compared the processing speed of Suite3D to a prior 3D analysis pipeline, the 3D version of CaImAn^15^, and found that it was substantially faster. To make the comparison, we considered a volume of 7 x 512 x 512 voxels, acquired at a rate of 3.3 Hz with a total of 400 frames with a conventional multi-plane twophoton microscope (one of the datasets used in Figure 3).

First, we compared the speed of motion correction. Suite3D performed the non-rigid motion correction on this dataset at a speed of 17 ± 1 volumes per second (mean ± s.d., n = 3 recordings). By comparison, the 3D variant of CaI-mAn^15,28^ processed only 0.9 ± 0.1 volumes per second (ED Figure 4).

Next, we compared the speed for detection and segmentation. We used standard cloud computing nodes for both Suite3D and the 3D mode of the CaImAn algorithm. We also used the faster online version OnACID^17^. Depending on the computing environment and processing parameters, Suite3D was typically 10-30x faster than CaImAn and 4-10x faster than OnACID (Figure 4k, ED Figure 4). Both versions of CaImAn require the user to estimate the number of cells before processing a movie (parameter K), and memory use scales with this parameter, making it challenging for large datasets (ED Figure 4). By contrast, the use of memory in Suite3D is independent of recording duration or number of cells, and is adjustable by varying batch sizes without affecting results.

Suite3D thus consistently provides substantial speedup in our experiments across all tested parameter configurations and datasets. Suite3D also allows users to sweep detection and segmentation parameters on subsets of a movie which allows users to quickly select the optimal parameters for their dataset, a process which requires custom scripts or many full re-runs in Suite2p, CaImAn or OnACID pipelines.

We further compared Suite3D to a powerful anatomical segmentation tool, CellPose^22^, which can segment 3D cell masks from a static 3D volume. As input to CellPose we provided the motion-corrected mean volume. CellPose missed almost half of the active cells in the volume (ED Figure 5), probably because many cells have sparse activity, and thus have a weak presence in the mean volume. Performance of CellPose also decayed in regions where neurons are tightly packed and difficult to unambiguously identify manually. Perhaps CellPose’s performance could be improved by using alternatives to the mean volume (e.g. the correlation volume), but due to the heterogeneity of neuronal time series, it is unlikely that a single metric would allow CellPose to capture all neurons.

**Figure 5.**
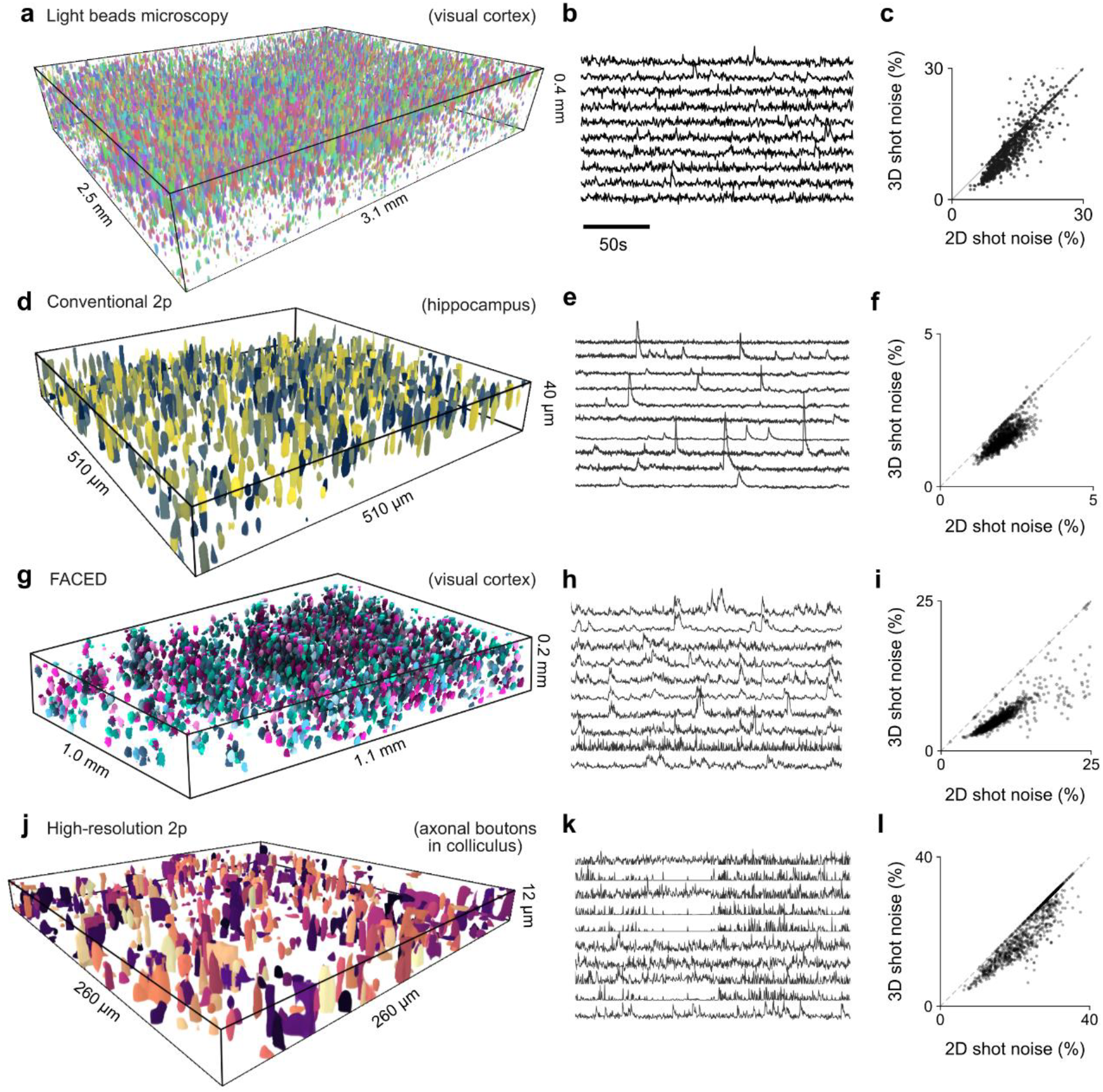
Application to diverse datasets. Suite3D works on volumes imaged across modalities, scales, noise levels and resolutions. **a**, The >40,000 neurons identified by Suite3D from a volume imaged with Light Beads Microscopy (LBM) in mouse visual cortex, with dense expression of GCaMP6s in excitatory neurons. **b**, Example fluorescence traces for 10 randomly selected ROIs from the volume in (a). **c**, Estimated shot noise for each 3D ROI and its corresponding 2D ROI acquired by considering only the voxels in the brightest plane for the ROI. **d-f**, Same, for a volume acquired with conventional multi-plane 2-photon imaging, with sequentially-acquired planes in mouse hippocampus with dense excitatory expression of GCaMP6s. **g-i**, Same for a volume imaged with FACED in mouse visual cortex with dense expression of GCaMP6s in excitatory neurons. **j-l**, Same, for a volume acquired with multi-plane 2-photon imaging at high resolution in mouse Superior Colliculus, showing boutons on retinal axons.

### Application to diverse datasets

Finally, we validated Suite3D over a range of imaging modalities, resolutions and brain regions (Table 1, ED Table 1), comparing its performance to that of a matched plane-by-plane analysis.

**Table 1.**
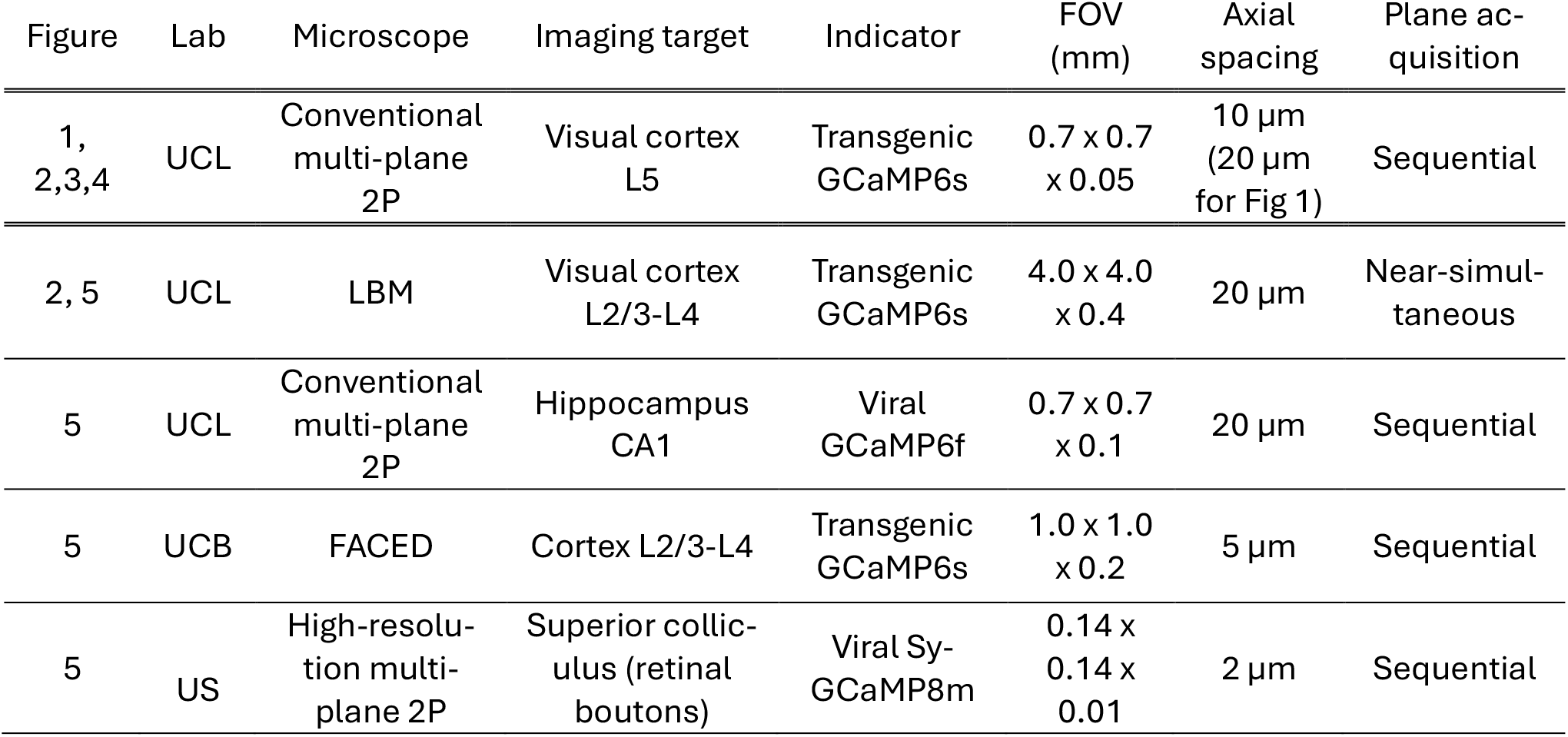
Datasets used in the development and testing of Suite3D.

| Figure | Lab | Microscope | Imaging target | Indicator | FOV (mm) | Axial spacing | Plane acquisition |
| --- | --- | --- | --- | --- | --- | --- | --- |
| 1, 2,3,4 | UCL | Conventional multi-plane 2P | Visual cortex L5 | Transgenic GCaMP6s | $0.7 \times 0.7 \times 0.05$ | $10 \mu\text{m}$ ($20 \mu\text{m}$ for Fig 1) | Sequential |
| 2, 5 | UCL | LBM | Visual cortex L2/3-L4 | Transgenic GCaMP6s | $4.0 \times 4.0 \times 0.4$ | $20 \mu\text{m}$ | Near-simultaneous |
| 5 | UCL | Conventional multi-plane 2P | Hippocampus CA1 | Viral GCaMP6f | $0.7 \times 0.7 \times 0.1$ | $20 \mu\text{m}$ | Sequential |
| 5 | UCB | FACED | Cortex L2/3-L4 | Transgenic GCaMP6s | $1.0 \times 1.0 \times 0.2$ | $5 \mu\text{m}$ | Sequential |
| 5 | US | High-resolution multi-plane 2P | Superior colliculus (retinal boutons) | Viral Sy-GCaMP8m | $0.14 \times 0.14 \times 0.01$ | $2 \mu\text{m}$ | Sequential |

We applied Suite3D to volumetric data where planes (separated by 20 µm) are acquired near-simultaneously with Light Beads Microscopy in the mouse visual cortex. Suite3D identified 43,652 neurons (Figure 5a). Traces from randomly selected example neurons show typical calcium transients (Figure 5b). Critically, for practically all neurons, the shot noise associated with the 3D ROI found by Suite3D was markedly lower than the corresponding 2D ROI (the largest single-plane slice of the same ROI, Figure 5c).

Similar results were seen for more common imaging configurations where planes are acquired sequentially, such as conventional multi-plane 2-photon imaging with a piezo-driven objective. In one such dataset (also with planes separated by 20 µm) obtained in mouse hippocampal area CA1, pyramidal neurons crossed multiple imaging layers (Figure 5d-f). This recording was acquired while the mouse navigated a virtual reality corridor, and the activity extracted by Suite3D reveals sequences of putative place cells (ED Figure 6).

We also applied Suite3D to volumetric recordings acquired with the FACED imaging method, which enables ultrafast line-scanning^4–6^ (Figure 5g-i). This recording was acquired during visual stimulation, and the activity extracted by Suite3D shows trial-locked responses to the visual stimuli (ED Figure 6).

To verify the performance of Suite3D on higher resolution recordings of non-somatic targets, we tested it on recordings of retinal boutons in superior colliculus (Figure 5j-l). In this recording the activity extracted by Suite3D shows a subset of boutons with synchronous global activity, while others carry asynchronous signals (ED Figure 6).

In all 4 test datasets Suite3D performed well with default parameters, but its performance was further optimized through built-in parameter tuning interfaces (ED Table 1). Indeed, as in all processing pipelines, the results can benefit from user-guided optimization of parameters. This process can be difficult in other pipelines: it may require re-running the entire dataset many times, writing code to extract intermediate results, or creating custom interfaces to visually compare segmentation results across tens of runs. Suite3D provides parameter sweeping interfaces that are integrated into the main processing pipeline. It can sweep tens of combinations of parameters on subsets of data in minutes in the detection and segmentation steps, and visualize intermediate results for the user to directly compare and gain an intuition of the parameters (ED Figure 7). A subset of parameters suggested for tweaking are presented in ED Table 2.

In combination, a robust set of default parameters and a nimble optimization interface make Suite3D well suited for efficiently analyzing volumetric data obtained with a variety of 2-photon imaging methods.

## Discussion

Volumetric datasets are increasingly common in 2-photon microscopy, and often lead to neurons appearing in multiple planes. As we have demonstrated with five example datasets, unless the planes are separated by distances that are substantially larger than the neurons, those neurons will appear in multiple planes. Indeed, depending on the target region and axial point spread function of the imaging system, the images of neurons can extend over 30 µm axially.

With such volumetric data, the current standard methods that process the data in 2D are inadequate. Typical 2D analysis pipelines can only correct for horizontal brain motion, leading to artifactual signals when motion in the axial direction leads to cells coming in and out of the imaging plane. Moreover, analyzing the data plane by plane leads to duplicated cells across planes, reduced signal-to-noise ratio per cell, and missed cells. Duplication could be reduced with manual or semi-automated curation^33,34^ but such approaches are not scalable to larger datasets.

By contrast, Suite3D is ideally positioned to process these volumetric data because it takes into consideration aspects of the data that are intrinsically 3-dimensional. First, it performs motion correction in 3D, which improves on noisy planar estimates of motion, and estimates not only horizontal motion but also axial motion. Second, it detects neurons using 3D correlation, which improves cell detectability. Finally, it performs 3D segmentation, detecting cells across imaging planes. The resulting cells have more voxels and therefore have lower levels of shot noise and stronger responses. At each processing step, therefore Suite3D improves over 2D methods largely because more voxels means better signal.

As confirmed by ground-truth data, Suite3D is also competitive with an existing 3D method of analysis. We obtained ground truth data with sparse anatomical labeling, and analyzed it using not only Suite3D but also the 3D version of an established pipeline, CaImAn^15^. Suite3D performed better than CaImAn in all metrics, including speed. A specific advantage of Suite3D is that its ROIs are compact and similar to the anatomical ground-truth, whereas CaImAn tends to find very large ones. Such large ROIs require demixing, which can remove useful signal^35^. A similar point has been made in the comparison between Suite2p and the 2D version of CaImAn^36^. However, demixing might be superior in situations where cells are tightly packed. In these circumstances, the neuropil correction used by Suite2p and Suite3D, which relies on the outer shell of a neuron, may not be optimal.

We have shown that Suite3D operates successfully on volumetric data from multiple microscopes, resolutions and brain regions. We considered data obtained with conventional multiplane 2-photon microscopy and with advanced volumetric 2-photon microscopes such as FACED^4–6^ and LBM^7,8^, which image multiple planes in quick succession. In all these datasets, Suite3D successfully detected cells appearing on multiple imaging planes, improving cell detectability and signal quality, and avoiding duplications. Suite3D might also be of use in analyzing volumetric data obtained with other types of microscope. For instance, there is a vast literature on imaging the whole brain of the larval zebrafish with light field^10,37–40^ or light sheet^41–46^ microscopy. These studies have typically used custom software packages and have not typically led to new general-purpose software packages for the analysis of volumetric data. It is possible that Suite3D would provide a good tool for analyzing those data.

The user experience of Suite3D is driven by three design principles: processing should be accurate, fast, and its steps should be transparent. Speed is an obvious concern when dealing with volumetric 2-photon microscopes, which can generate close to 1 TB/hour, but it also has a less obvious benefit: faster algorithms allow advanced users to tinker and optimize parameters on challenging datasets. By running quickly, Suite3D allows users to iterate. Crucially, a cell detection pipeline should be both efficient and accurate^47^: Suite3D’s speed does not come at the cost of detection quality.

Suite3D is thus a useful tool for volumetric imaging applicable across a variety of microscopes and scientific questions. We released it to the community as an open-source package and welcome suggestions and contributions to further improve it.

## Acknowledgments

We thank Carsen Stringer and Marius Pachitariu for invaluable suggestions at the beginning of this project and their continued development of Suite2p. We thank Bex Terry, Kevin Barber, Francisca Martínez Traub, and the UCL and Rockefeller University animal facilities for animal care and management; Tobias Nöbauer and Jeffrey Demas for advising microscope development; Flynn O’Connell for contributions to the software; Santiago Otero Colonel, Paul Fahey and Max Gagnon for testing the software; and Kimberly Ren for feedback on the manuscript and figures.

This work was funded by UKRI (Frontier Award EP/X022366/1 to MC), BBSRC (grant BB/W019884/1 to MC), the National Institutes of Health BRAIN initiative (grant U01NS126057 to AV and MC), the Wellcome Trust (Investigator Award 223144/Z/21/Z to MC and KDH), and the ERC (101097874 to KDH), the National Institutes of Health BRAIN initiative (UF1NS107696, U01NS118300, U01NS137449 to NJ and JZ), and jointly by the Wellcome Trust and the Royal Society (Sir Henry Dale Fellowship 220169/Z/20/Z to SS). AH was supported by a SWC student fellowship (Gatsby Charitable Foundation GAT3755 and the Wellcome Trust 219627/Z/19/Z), by a UCL Bogue travel fellowship, and by the National Institute of Diabetes and Digestive and Kidney Diseases (DP1DK139958 to Mark Andermann). 3D visualizations were performed with UCSF Chimera, developed by the Resource for Biocomputing, Visualization, and Informatics at the University of California, San Francisco, with support from NIH P41-GM103311. MC holds the GlaxoSmithKline / Fight for Sight Chair in Visual Neuroscience.

## Data Availability

The five example datasets used in this manuscript are listed in Table 1 and are available via FigShare (https://doi.org/10.5522/04/32956220).

## Code Availability

Suite3D is released as an open-source software package and can be found at https://github.com/alihaydaroglu/suite3d/. Code to analyze the five example datasets can be found in the FigShare repository (https://doi.org/10.5522/04/32956220). Documentation, demos, and interactive visualizers can be found at https://suite3d.github.io.

## Author contributions

According to the [CRediT](https://credit.niso.org/) taxonomy.

|  | Agarwal | Baruchin | Carandini | Chang | Cozan | Dodgson | Guo | Harris | Haydaroglu | Ji | Krumin | Landau | Meyer | Reddy | Schröder | Vaziri | Zhong |
| --- | --- | --- | --- | --- | --- | --- | --- | --- | --- | --- | --- | --- | --- | --- | --- | --- | --- |
| Conceptualization |  |  | ● |  |  |  |  | ● | ● |  |  |  |  |  |  |  |  |
| Data curation |  |  |  |  |  |  |  |  | ● |  |  |  |  |  |  |  |  |
| Formal analysis |  |  |  |  |  |  |  |  | ● |  |  |  |  |  |  |  |  |
| Funding acquisition |  |  | ● |  |  |  |  | ● |  | ● |  |  |  |  | ● | ● |  |
| Investigation |  | ● |  | ● | ● | ● | ● |  | ● |  | ● | ● | ● | ● |  |  | ● |
| Methodology |  |  | ● |  |  |  |  | ● | ● |  | ● |  |  |  |  | ● |  |
| Project administration |  |  | ● |  |  |  |  |  |  |  |  |  |  |  |  |  |  |
| Software | ● |  |  |  |  | ● |  |  | ● |  |  | ● |  |  |  |  |  |
| Resources |  |  | ● |  |  |  |  | ● |  | ● | ● |  |  | ● | ● | ● |  |
| Supervision |  |  | ● |  |  |  |  | ● | ● | ● |  |  |  |  | ● | ● |  |
| Validation |  |  |  | ● |  |  |  |  | ● |  |  | ● |  |  |  |  |  |
| Visualization |  |  | ● | ● |  |  |  |  | ● |  |  |  |  |  |  |  |  |
| Writing – original draft |  |  | ● |  |  |  |  |  | ● |  |  |  |  |  |  |  |  |
| Writing – review & editing |  |  | ● |  |  |  |  | ● | ● | ● |  |  |  |  | ● | ● |  |

## Methods

Here, we first describe the processing steps in Suite3D. We then provide brief methods for the acquisition of the data that are analyzed in this study, and for the comparisons between analysis pipelines.

### Suite3D

Suite3D is written in Python and is available under a GNU Affero GPL v3.0 open-source license (www.gnu.org/licenses/agpl-3.0.en.html). Below we describe its processing steps, and we provide pseudocode.

#### Preprocessing

Large-FOV 2-photon microscopes often use a resonant-galvo-galvo (RGG) scanning configuration for extended lateral scanning^5,48^. This results in the field of view being split into “strips”, which often overlap at their edges. Suite3D uses the cross-correlation between edges of imaging strips to find the optimal shift between strips before fusing them together to construct a single continuous image.

Temporally multiplexed imaging methods such as Light Beads Microscopy can introduce inter-plane crosstalk since the inter-voxel time interval is similar to the fluorescence half-life of calcium indicators. We assume the following unidirectional model of crosstalk, where *F*_*i*_ is the true fluorescence and *S*_*i*_ is the measured signal in plane *i*, and *α* is a scalar “crosstalk coefficient”:

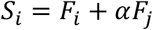

For a user provided set of cross-talking pairs {(*i*_1_, *j*_1_), … (*i*_*P*_, *j*_*P*_)}, the default algorithm in Suite3D finds *α* that minimizes second derivative of the correlation coefficient with respect to the crosstalk coefficient, across all pairs:

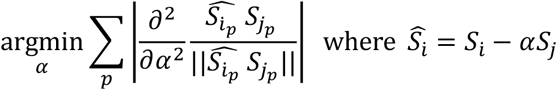

We find that this approach outperforms minimizing the correlation coefficient directly, both in simulated and real data (ED Figure 8). This is due to similarities in low-spatial frequency structural features (blood vessels, neuropil) which create a non-zero correlation coefficient between the true signal in any pair of planes (*F*_*i*_, *F*_*j*_). Minimizing the correlation coefficient directly causes an overestimate of the crosstalk coefficient. Our method finds a transition point in the crosstalk subtraction, which corresponds to a point where content corresponding to cells is fully removed and the remaining correlation is due to shared anatomy. Alternatively, the user can set a parameter to minimize the mutual information between *S*_*i*_, *S*_*j*_, similar to the PICASSO method^49^.

The final step of the preprocessing phase is the construction of a “reference volume”. Often, volumetric data is acquired as a series of axially separated planes between which there may be constant lateral offsets due to optical considerations. Suite3D uses a subset of the recording (typically 200-400 frames) to register planes to one another and undo inter-plane lateral offsets. Then, an iterative version of the volumetric motion correction algorithm (described in the next section) is applied to the subset of frames to construct a reference volume. This reference volume is often “sharper” than a simple time-averaged version of the movie because the brain motion has been corrected, so it is later used as a template during motion correction.

In practice, the computation of the preprocessing parameters is merged into a single “initialization” step which computes the optimal fusing and crosstalk values, constructs a reference volume and computes several quality metrics of the recording within several minutes on a subset of the full movie. In a typical workflow, this serves as a checkpoint where the user has the option to use the GUI to verify the recording quality, exclude noisy planes and confirm processing parameters before moving onto the more time-consuming stages. In future processing steps, whenever a batch of frames is loaded from disk, the described preprocessing steps (fusing, crosstalk subtraction, plane-to-plane registration) are applied with precomputed parameters as part of the I/O pipeline.

#### 3D motion correction

Rigid volumetric motion-correction is implemented with a phase-correlation based algorithm, similar to one previously described in Ref. ^13^. We replaced the Fast Fourier Transform (FFT) step with a three-dimensional version and extended the multiplicative and additive spatially tapered masks to three dimensions to reduce artifacts at the edges of the volume. The resulting 3D phase correlations can be interpreted similarly to 2D phase correlation maps, where the location of the peak indicates the three-dimensional displacement that must be applied to the given frame. The same method is first applied to the full rigid volume, and then to overlapping 3D sub-volumes (“blocks”). To estimate the location of the peak at subpixel resolution, we use kriging^13,28^. Rigid, sub-pixel translations are applied with trilinear interpolation. To apply non-rigid translations, which assign a x, y and z shift for each block, we construct a smooth 3D field of shifts over the entire volume, and correct them using trilinear interpolation^13^. We found that in higher noise, lower-resolution recordings, where applying 2D non-rigid correction can be detrimental to the data quality, 3D correction can still correct non-rigid motion.

To accelerate motion correction, we developed a novel GPU-based implementation of the rigid and nonrigid motion correction algorithms. In our experiments, naïve porting of the Suite2p motion correction algorithm to GPUs did not yield speed improvements, due to the costly nature of transferring large image data between RAM and GPU VRAM. To minimize the back-and-forth data transfers, we implement rigid and non-rigid registration steps directly on the GPU using a combination of CuPy-accelerated Python code and custom CUDA kernels in C++.

For a typical use case of Suite3D in a laboratory environment with active acquisition of large-scale volumetric data, one of the major bottlenecks is the file transfer speed. The raw data is typically moved from the experiment workstation to a shared server, from where it is typically moved to a fast disk on a processing workstation. Running pipelines such as Suite2p or CaImAn on data that is located on a server can cause significant slowdown, depending on I/O speeds. Suite3D motion correction implements a multi-threaded file I/O pipeline, where the next batch is loaded from storage into memory while the current batch is processed on the GPU, thus removing the requirement of moving data to local storage and masking the I/O times. Nevertheless, data I/O remains the biggest speed bottleneck, and ensuring fast read and write of data to a local SSD typically results in substantial improvements in throughput.

#### 3D detection

The correlation volume is inspired by the 2D correlation map implemented in Suite2p. Before the motioncorrected movie is passed to the algorithm, it is temporally binned – the size of the temporal bin is an optional user parameter that should be set to the number of frames in a typical transient (*detection_timebin*). If not provided, it is inferred from the time constant of the indicator. Next, the binned movie is temporally high-pass filtered with a rolling mean filter with a ~10s window to exclude low-temporal frequency variations. Next, each pixel is normalized by the smoothness of its temporal activity, estimated by the standard deviation of its first temporal differences to account for varying signal-to-noise levels. To remove the neuropil-driven shared variability across pixels, the movie is spatially high-pass filtered with a cutoff around 70 µm to create the detection movie.

Next, the detection movie is spatially low-pass filtered to enhance structure at the functional length scales. These filters can be changed depending on the targets. For example, when imaging axonal boutons, the cutoff for the functional filter should be set to <1 µm. On this filtered movie, the root-sum-square of each pixel across time is computed, only considering frames where the value of the given pixel was above the activity threshold, reducing the movie across the time dimension. The resulting correlation volume is then thresholded and used as the seed for cell segmentation. Suite3D allows the user to sweep a range of values for the sizes of the filters, the value of the threshold, and the strength of the normalization on a subset of the movie quickly, to manually optimize over the default settings if necessary for a particular dataset.

Suite3D implements several substantial algorithmic changes relative to Suite2p. First, the 2D spatial filters for neuropil subtraction and cell detection are extended to 3D. This allows improved detection for cells extending over several planes, and better estimation of neuropil at each voxel. The user has the option to select between Gaussian and top-hat filters for each. Second, Suite3D implements a local thresholding method^50^ which adaptively thresholds the correlation map at each location. This approach is beneficial when imaging large field-of-views or using microscope configurations that may lead to uneven SNR at different imaging depths, where the correlation map will scale differently for each region of the imaged volume. This can be problematic for the segmentation step, as a single hard threshold across the full volume will be too low for some regions and too high for others. Third, Suite3D introduces an adjustable pixel normalization parameter that scales the strength of normalization. This makes the temporal-difference normalization more robust. Otherwise, in high-noise recordings, dark regions (such as blood vessels) can become bright in the correlation map. This artifact is remedied by setting the normalization parameter to 0.7-0.9. Finally, Suite3D introduces the explicit on-disk detection movie, which disentangles the correlation map computation from cell segmentation. Particularly for noisy and low-resolution dataset, this allows much easier tuning of parameters.

The memory and processing requirements of computing the correlation map in one go as implemented in Suite2p are often unfeasible for large volumetric recordings. Suite3D, therefore, computes the correlation map with a batch-wise algorithm. For running estimation of standard deviation across batches, we used Welford’s algorithm^51^. The filtering and reduction steps are parallelized across CPU cores. To handle large arrays which cannot be held in memory at once, Suite3D uses efficient memory-mapped, parallelized algorithms supported by Dask (docs.dask.org/en/stable/) and Zarr (zarr.readthedocs.io/en/stable/).

##### Algorithm 1

Batch-wise Correlation Map Computation

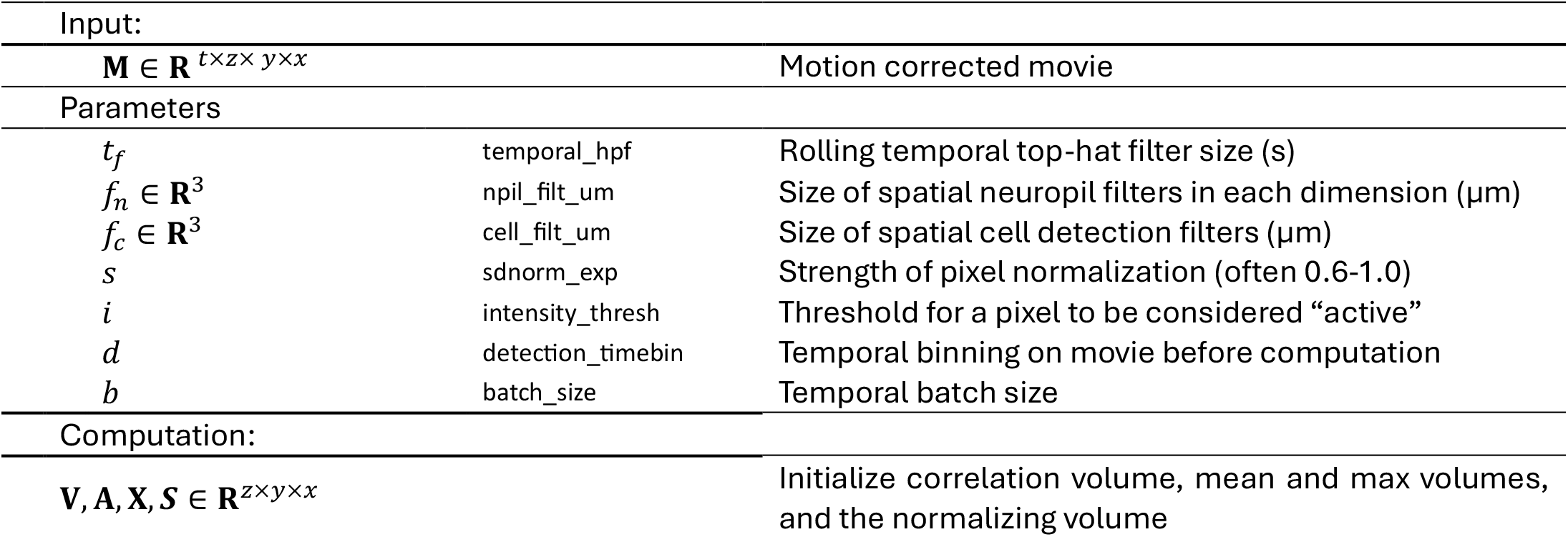
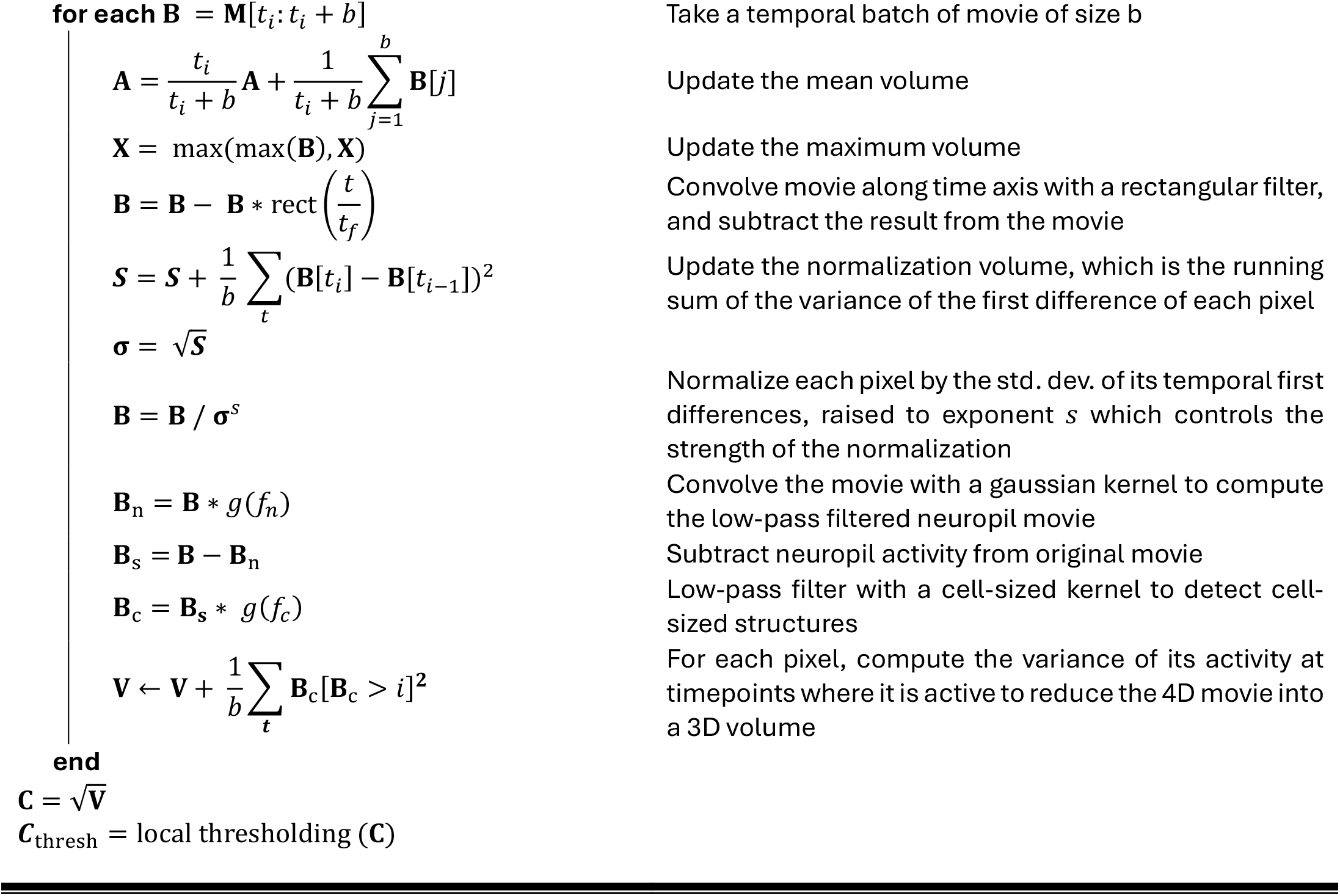

#### 3D segmentation

The segmentation phase iteratively selects candidate voxels from the correlation volume and expands them into ROI masks based on the correlation of their activity with neighbors. First, a seed voxel is selected from the correlation volume by taking the maximum and initializing a seed region of interest (ROI) with 9 voxels. Next, the neighboring voxels are taken as candidate voxels for expanding this ROI. The candidate pixel selection and growth of the ROI happens in three dimensions, allowing Suite3D to compute volumetric cell masks. The activity of each candidate voxel is compared to the activity of the ROI in the neuropil subtracted movie saved in the detection step. The comparison uses only frames where the existing ROI is considered “active”, by thresholding its activity by a hard threshold or 99^th^ percentile cutoff, whichever is lower (both parameters are user-adjustable). Suite3D computes a voxel-SNR metric for each candidate voxel and the ROI and compares it to a set threshold to determine whether a candidate voxel is included.

The voxel-SNR is computed by projecting the active frames of the voxel onto the unit-normalized ROI activity vector to compute the variance explained by the ROI activity on the voxel. Next, we compute the ratio of this variance to the non-ROI-related variance of the voxel to compute a signal-to-noise fraction (not to be confused with the shot noise fraction used in Figure 4,5). Each included voxel is weighted by its projection on the ROI activity. We find that this algorithm is more robust to noise than the method implemented in Suite2p, and that it improves the segmentation results. Suite3D also provides the user an option to revert to the Suite2p-style method where the mean over active frames is used to determine expansion. This expansion step is executed iteratively (see Algorithm 2) until an ROI cannot be expanded anymore, after which a new seed voxel is selected for the next ROI.

The Suite2p algorithm can fail in lower-resolution datasets by detecting “junk” ROIs that are too small, too large, or too disjointed to be real cells. Suite3D re-casts the algorithm to create several interpretable parameters, which are exposed through a parameter selection UI and can be adjusted to solve the problems. *c*_min_ controls the number of ROIs detected by changing the endpoint of the segmentation step – if obvious ROIs are being missed, this parameter should be lowered. *e*_thr_ controls how eager the algorithm is to expand a given ROI: if ROIs are too small, this should be lowered, if ROIs extend too wide, this should be increased. *p*_active_ and *i*_thr_ jointly determine the active frames of each ROI – for short recordings, lowering the thresholds tend to give better results, while for longer recordings (~hours) the thresholds can be set very high so only high-quality timepoints are considered as “active” frames.

##### Algorithm 2

Parallelized volumetric cell segmentation

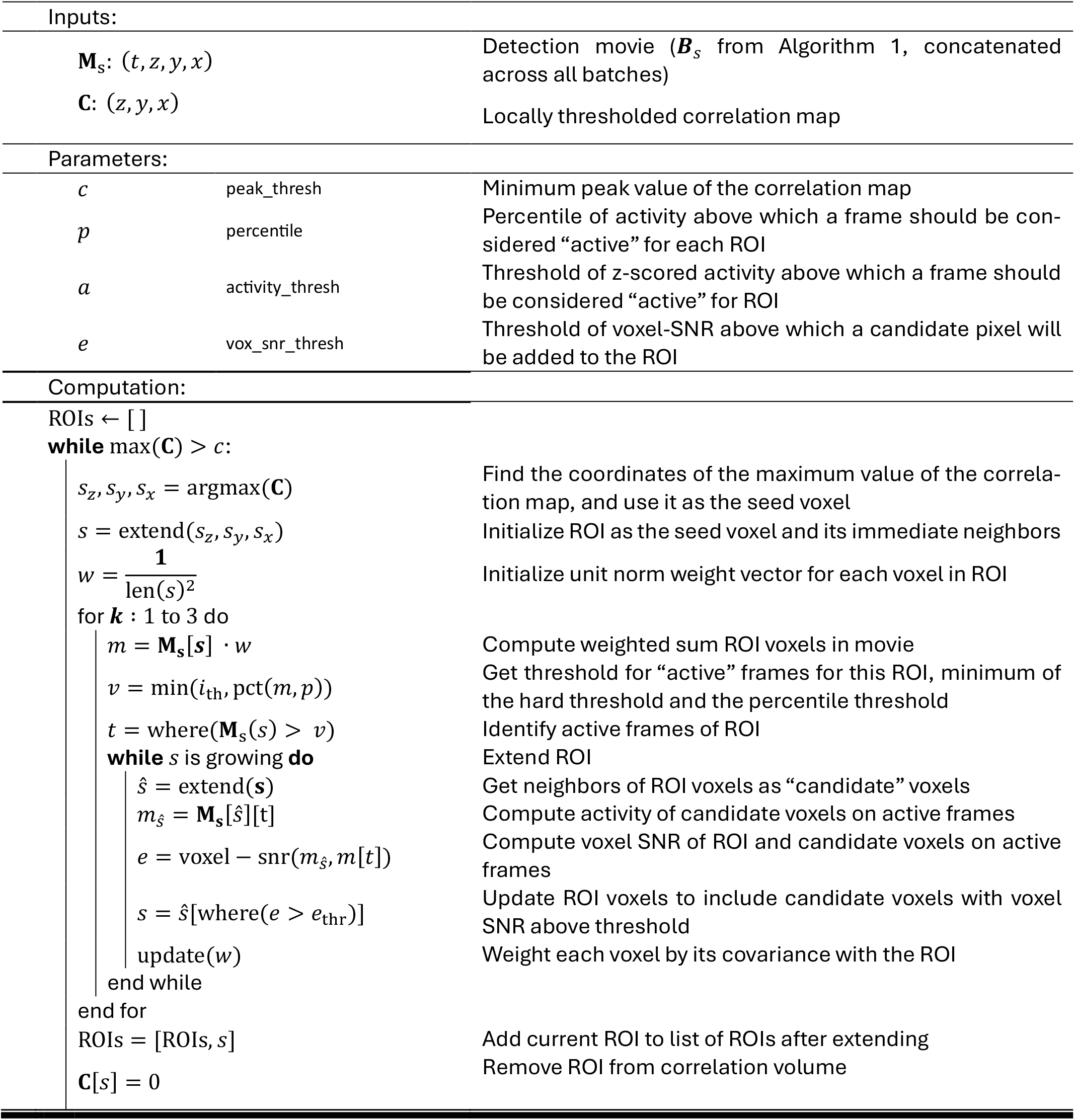

Computationally, the segmentation algorithm is demanding as it requires the entire time series of the movie to be held in memory at once. To make it tractable, the movie is split into patches in the x and y dimensions, and each patch is loaded entirely into memory one at a time. The detection movie used is the one computed in the correlation map computation step, which is located in a temporary file in fast local storage and accessed through memory-mapping. To account for cells that may cross boundaries between patches, small overlaps between patches are enforced and cells are de-duplicated at the end of the segmentation step. For a given spatial batch, the algorithm described is not straightforward to parallelize since the segmentation of each cell requires updating the correlation map, which is then required to identify the next cell to segment. To parallelize across *n* CPU cores, our algorithm greedily selects the top *n* peaks of the correlation map with a minimum distance constraint between all pairs of selected peaks. Then, each worker accesses the subtracted movie located in shared memory to segment, and each updates the correlation map in shared memory once their selected cell is extracted. This is implemented using the multiprocessing module, and specifically the SharedMemory class in Python.

#### Postprocessing and curation

Once weighted masks are computed for each region of interest, Suite3D returns to the motion-corrected movie to extract the weighted average of fluorescence over all pixels in each ROI. In addition, Suite3D implements a novel volumetric neuropil correction – a 3D hollow shell around each ROI is computed (excluding other ROIs), and the average activity within this hollow shell is taken to be the neuropil fluorescence. This approach is a volumetric adaptation of the Suite2p neuropil correction. This neuropil fluorescence is subtracted from the ROI fluorescence for each ROI with a coefficient of 0.7. Next, the neuropil-subtracted traces are deconvolved using a version of the OASIS algorithm with no L0/L1 constraints^30,31^.

Once ROIs are extracted, a custom GUI based on napari^52^ is used for semi-automated curation and quality control. The interactive GUI allows histogram-based filtering of ROIs based on size, values on the correlation volume, or activity levels, and is easily extensible to include user-defined cell filters. It also allows manual curation of individual ROIs, allowing the user to mark them as cells and non-cells. For high-quality 3D visualization, Suite3D exports cell maps in the .mrc format for visualization in Chimera^53^, which was used for visualizations in Figure 5.

### Dataset Acquisition

To test and validate Suite3D, we acquired datasets at University College London, and we analyzed datasets previously obtained at Rockefeller University, University of California Berkeley, and University of Sussex.

#### University College London

Experiments at University College London (UCL) were conducted according to the UK Animals Scientific Procedures Act (1986) under personal and project licenses released by the Home Office following appropriate ethics review. Experiments in L2/3 of visual cortex were performed on 5 adult mice (3 female, 2 male; aged 11-25 weeks) expressing GCaMP6s in all cortical excitatory neurons (CaMKII x Ai162). Experiments on Layer 5 of visual cortex with sparse anatomical labelling were performed on an adult female mouse (aged ~20 weeks) expressing Cre recombinase GCaMP6s in excitatory neurons of cortical layer 5 (Rbp4-Cre x Ai94). Ultra-sparse structural labelling was achieved via intracranial injection of adeno-associated viruses carrying Cre-dependent Flp recombinase (AAV-EF1a-DIO-FLPo-WPRE-hGHpA) and Flp-dependent tdTomato (AAV-hSyn-(ATG-out)-fDIO-NLS-tdTomato-E2A-N2cG). Experiments on the CA1 region of the hippocampus were performed on adult mice aged 8-20 weeks expressing GCaMP6f in excitatory neurons (AAV9-CaMKII-GCaMP6f).

Mice were implanted with cranial windows of 4 mm diameter under surgical anesthesia. Details of the surgery can be found in Ref. ^54^. For experiments on CA1, we aspirated the cortical tissue above the hippocampus and implanted a cylindrical imaging cannula with a coverslip sealing the lower opening^55^. After recovery (minimum 5 days), mice were head-fixed and free to run on a low-resistance treadmill during imaging. In some recordings, visual stimulation was delivered via three screens covering approximately 270° horizontally and 70° vertically of the visual field of the animal. Full-field drifting gratings were presented with 12 directions (30° spacing) in random order, with eight repeats of each unique stimulus. Each grating was presented for 2 s, with a 2-3 s inter-stimulus interval. In CA1 recordings, mice were performing a virtual reality task involving running through a linear corridor presented on the same screens, while head-fixed on a treadmill.

Light Beads Microscopy (LBM) recordings at UCL were conducted with a modified Many-fold Axial Multiplexing Module (MAxiMuM) cavity^7^, integrated with a commercial mesoscope (Thorlabs, Multiphoton Mesoscope). The laser source comprised a 4.8 MHz pump laser (Coherent Monaco, 60W average power at 1030 nm) with an optical parametric chirped-pulse amplifier (OPCPA, White Dwarf from Class 5 Photonics), with a maximum average output power of 6W at 960 nm.

Conventional multi-plane imaging was conducted on a commercial 2-photon microscope (Bergamo II, Thorlabs). Sequential acquisition of planes was achieved by piezo-driven actuation of the objective. Visual cortex imaging was conducted on transgenic mice expressing GCaMP6s in Layer 5 excitatory neurons (Rbp4 x Ai162), with similar cranial windows as described above. Hippocampal imaging was conducted on mice with viral expression of GCaMP6f in CA1, imaged through an optical cannula.

#### Rockefeller University

LBM recordings at Rockefeller University were conducted in accordance with protocols approved by the Institutional Animal Care and Use Committee (IACUC) on adult transgenic mice (2 female; aged 10-20 weeks) expressing GCaMP6s in excitatory neurons (vGlut1-Cre x Ai162). The imaging was performed on a similar LBM as described above, using similar methods.

#### University of California Berkeley

FACED microscopy at University of California Berkeley was conducted in accordance with protocols approved by the Institutional Animal Care and Use Committee (IACUC). The dataset that was analyzed was obtained in a mouse expressing GCaMP6s in excitatory neurons (Slc17a7-IRES2-Cre × Ai162D). The imaging was done with a lateral voxel spacing of 1.4 µm and 2.0 µm in x and y, and an axial spacing of 5 µm in z. Further details can be found in Refs. ^4,6^.

#### University of Sussex

High-resolution imaging of retinal boutons at the University of Sussex was conducted according to the UK Animals Scientific Procedures Act (1986) under personal and project licenses released by the Home Office following appropriate ethics review. The dataset that was analyzed was obtained in a mouse expressing SyGCaMP8m virally^56^. A conventional 2-photon microscope was used, with lateral voxel spacing of 0.5 µm and an axial spacing of 2 µm on mice. Further details can be found in Ref. ^33^.

### Validation

#### Anatomical Ground Truth

We used recordings with sparse anatomical red labelling with tdTomato as our source of “ground truth” labels. In these recordings, red and green channels were simultaneously acquired during visual stimulation. The red channel was sufficiently bright that no green crosstalk was visible due to the high relative brightness of the red indicator.

We first computed a reference volume on the red channel of recordings using Suite3D, to correct for motion. Then, we co-aligned the red reference volume to the reference volume from functional recordings using per-plane translation. We used recordings with no substantial z-drift measured in the green channel. We clipped and normalized the red volume to its 99.5^th^ percentile, and ran CellPose 3D on this volume. We swept the flow_threshold (0.4 – 1.0), cellprob_threshold (0.0 – 3.0) and diameter (12-20 and “auto”) parameters and visually inspected the results to find the best performing segmentation of the red volume. The optimal combination set flow_threshold at 1.0, cellprob_threshold at 1.0 and diameter at “auto”. The resulting masks are shown in ED Figure 9.

We next matched functionally-detected ROIs with the anatomical ground truth ROIs to evaluate performance. First, we removed from our analysis anatomically-detected cells that were silent in the recording. For each anatomical ROI, we extracted its neuropil-corrected activity by regressing out a surrounding neuropil mask from the mean activity of voxels within the mask. We computed the ratio of the standard deviation of the within-mask activity with that of a mask of surrounding neuropil. We selected a ratio of 1.3 as the silence threshold based on manual inspection of traces. In other words, the activity of a cell must have at least 30% larger standard deviation than surrounding neuropil to be considered active. Of the 200 anatomical ROIs detected, 73 passed this threshold. The large number of silent cells is possibly due to the short recording duration (<20 minutes).

We then loaded 3D ROI masks, detected either by Suite3D (“lam” vectors from stats.npy) or 3D CaImAn (“estimates.A”). For per-ROI figures (i.e. Figure 3n), we used the configuration of CaImAn that found the most similar number of ROIs to the corresponding Suite3D run on the same dataset. For Figure 3o,p, we compute metrics on ROIs from all 36 configurations of CaImAn we used. Further details on CaImAn implementations are discussed in the next subsection. For each pair of functional and red ROIs, we computed the cosine similarity of spatial masks such that fully overlapping ROIs would score 1 and non-overlapping ones score 0. We considered a red cell “matched” with its maximum-overlap functional counterpart if the overlap exceeded 0.3. We then computed the activity correlation of matched pairs computing the correlation coefficient between the high-pass-filtered ground truth trace and the functionally-extracted trace. In the case of Suite3D, this corresponds to the neuropil subtracted trace, whereas for CaImAn we used the denoised trace.

To evaluate to what extent red cells were uniquely detected by pipelines, we computed the “recall” and “uniqueness” of detected cells. We define recall as the fraction of active red cells that had at least one functional ROI with spatial similarity greater than 0.3. We did not compute precision directly, as there were many more green cells than red cells due to the sparse expression strategy, and these would wrongly be classified as false positives. Instead, we computed the uniqueness of each red cell’s matches as 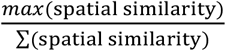. A red cell that overlaps with only a single green cell would have a score of 1, and one that overlaps equally with two cells would have a score of 0.5. We found that many CaImAn ROIs had very large footprints with relatively low weights outside of a cell-sized core, which reduced the uniqueness score as a single red cell was often overlapped by many low-weight tails of other ROIs. To avoid this confound, we clipped both functional ROIs to only include voxels that sum to 75% of the total weight of the full ROI. This removed the extended tails from CaImAn. We repeated the analyses in Figure 3 varying the ROI fraction (50% - 100%) and the spatial similarity threshold (0.1 – 0.4), which did not change the conclusions.

#### Functional Validation

To evaluate the functional signals extracted with Suite3D and compare it to 2D pipelines, we analyzed responses to drifting gratings. We used data acquired from V1 with a conventional multi-plane 2p microscope, and analyzed it with default Suite3D parameters (ED Table 1, row 1). We also detected ROIs, running Suite2p on each plane.

To match cells detected in Suite2p and Suite3D, we used a semi-automated method considering the distance between the centroid of each pair of cells across modalities, and the correlation of their extracted fluorescence. To ensure that matches were genuine, we chose conservative thresholds for correlation and distance. Then, we estimated fraction of shot noise^32^ for the extracted fluorescence traces for matched ROIs across cells.

To compare variability across trials, we normalized each fluorescence trace by its standard deviation and extracted responses in the time windows where stimuli were presented. We took the standard deviation of each ROI’s fluorescence across eight repeats of the same stimulus, averaged over all unique stimuli.

To compute the number of significantly stimulus-responsive cells in each pipeline, we considered all ROIs (matched and non-matched). For each ROI, we took the average activity in the time period from 0.5s after stimulus onset to 2.0s after stimulus onset. We computed the fraction of signal-related variance across two halves of repeats of each stimulus as described in ref^54^. To identify statistically significant cells, we repeated the same computation 500 times, shuffling the stimulus identities. Cells with a signal-related variance fraction above the 99^th^ percentile of the null distribution were considered significantly responsive.

We then investigated the duplication and omission of cells with 2D pipelines. For both Suite2p and Suite3D ROIs from conventional V1 datasets, we computed the pairwise correlations of the deconvolved activity for all pairs of ROIs whose centroids fell within 20 µm in x/y. We compared the distribution of this duplication score for Suite3D and Suite2p ROIs (Figure 4d). To further characterize the difference between 2D and 3D segmentation we ran Suite3D in “2D mode” by disabling volumetric features. This allowed us to isolate the benefit of moving from 2D to 3D segmentation. On the same dataset, we assigned each 2D ROI to a 3D ROI based on pixel overlap. We quantified how many 2D ROIs each 3D ROI got assigned to, and how many 3D ROIs were omitted by the 2D version (ED Figure 3). We further quantified the pairwise correlation distribution for pairs of 2D ROIs belonging to the same 3D ROI. This is relevant for post-hoc merging algorithms, which rely on finding a hard threshold between this distribution and the distribution of 2D ROI pairs from different 3D ROIs.

### Hardware and Software

#### Computing environments

All experiments in the paper were conducted on one of three computing environments: a local workstation, an 8-CPU cloud instance (r6a), or a GPU-equipped cloud instance (g4dn).

The local workstation was equipped with an Intel Xeon w9-3475 CPU with 36 physical cores (72 threads) operating at a base frequency of 2.2 GHz, with 512 GB of DDR5 RAM, and an NVIDIA RTX A4500 GPU with 20 GB of VRAM. The workstation was equipped with a pair of 4TB M.2 SSDs in RAID0 configuration and had a 10 Gbps connection to the server on which raw data was stored. This workstation was used to generate all results in main figures, and some of the benchmarking of all pipelines in ED Figure 4.

The GPU-equipped g4dn cloud instance was used to benchmark Suite3D. It is provided through Amazon Web Services (AWS) Elastic Compute Cloud (EC2), with the identifier g4dn.2xlarge. It is equipped with 8 vCPUs (Cascade Lake), 32 GB of RAM, and an NVIDIA Tesla T4 with 16 GB of VRAM and a local NVMe. We used the Deep Learning Base OSS NVIDIA Driver GPU AMI (Ubuntu 22.04). The local NVMe was used to store raw data and intermediate files.

The r6a cloud instance was used to benchmark CaImAn and OnACID. It is provided through AWS EC2 with identifier r6a.2xlarge. It is equipped with 8 vCPUs (AMD Zen 3), 64 GB of RAM, and the AWS Elastic Block Store tier 3 general purpose SSD (gp3). We used the Ubuntu 22.04 AMI. The gp3 store was used to store raw data and intermediate files. The AMD Zen 3 vCPUs are a newer generation than the Cascade Lake vCPU of the g4dn instance.

#### CaImAn

We used CaImAn 1.12.2 for all comparisons, both on the local workstation and on the r6a cloud instance. We evaluated CaImAn on the V1 dataset acquired with conventional multi-plane 2p imaging used in Figures 3 and 4. The parameters and workflow from the “demo_caiman_cnmf_3D.ipynb” notebook from the CaImAn repository was used. Parameter “gSig” was set to (5,5,2) from the default of (4,4,2) and “fr” was set to 4 Hz to match our spatiotemporal recording parameters. We then swept the values of three parameters, varying “K” (expected # of components) between (100, 500, 1000, 5000); “merge_thresh” between (0.4, 0.8, 0.9) and rval_thr between (0.3, 0.6, 0.9). When benchmarking CaImAn on cloud instances, we ran into RAM issues on the g4dn instance used for Suite3D, so we moved to the r6a instance with larger capacity. As in the demo file, we used a fresh worker cluster for each parameter combination, ran an initial fit, followed by component evaluation and refitting. We then aligned the CaImAn outputs to Suite3D outputs via rigid shifts, and converted the CaImAn components and traces into the Suite3D stats.npy format for direct comparison.

We also evaluated OnACID, CaImAn’s online-CNMF implementation, which is packaged within the 1.12.2 installation. Our OnACID pipeline followed approach 1 of the demo_realtime_cnmfE.ipynb notebook, with multi-frame parallelism. We used the same parameters as the offline CaImAn configuration, but had to set p=0 and skip deconvolution due to errors. We swept the value of “K” between (200, 500, 1000) and min_SNR between (2,3,4) with rval_thr=0.9.

During runs, we ensured both OnACID and CaImAn had high utilization of their allocated CPUs, to ensure they were not bottlenecked by file I/O or memory. In some cases, reducing the CPU allocation to 16 from 36 increased CaImAn’s runtime – in these cases, we used the reduced allocation for benchmarking. In benchmarking experiments, datasets fit in RAM and hence were not bottlenecked by memory mapping. The Dockerfile used to run the AWS benchmarking can be found in the Suite3D repository, under “suite3d/benchmarking/paper/Dockerfile.caiman”, alongside scripts used to benchmark Suite3D, CaImAn and OnACID on AWS.

#### Suite2p

We used version 0.14.4 and 1.1.0 of Suite2p, as implemented in github.com/MouseLand/Suite2p. We found no substantial difference in results between two versions. We ran the processing independently on each plane with default parameters, and swept the threshold_scaling parameter between (1.0, 2.0, 2.5, 3.0, 4.0, 5.0).

#### CellPose 3D

We used version 4.0.7 with default cpsam weights. In addition to its use during ground-truth validation, we also tested CellPose 3D as an alternative to functional segmentation by running anatomical segmentation on a static image. We passed the motion corrected Suite3D reference image of the green functional channel from a conventional V1 dataset to CellPose. We swept the flow_threshold over (0.05, 0.1, 0.2, 0.25, 1.0, 2.0) and cellprob_threshold over (−5, −3, −2, 0) and selected 0.2 and −2 respectively. We converted the resulting integer-label mask volumes to the Suite3D format, and used the Suite3D neuropil subtraction and extraction pipeline (ED Figure 5).

## Extended Data Figures and Tables

**ED Figure 1.**
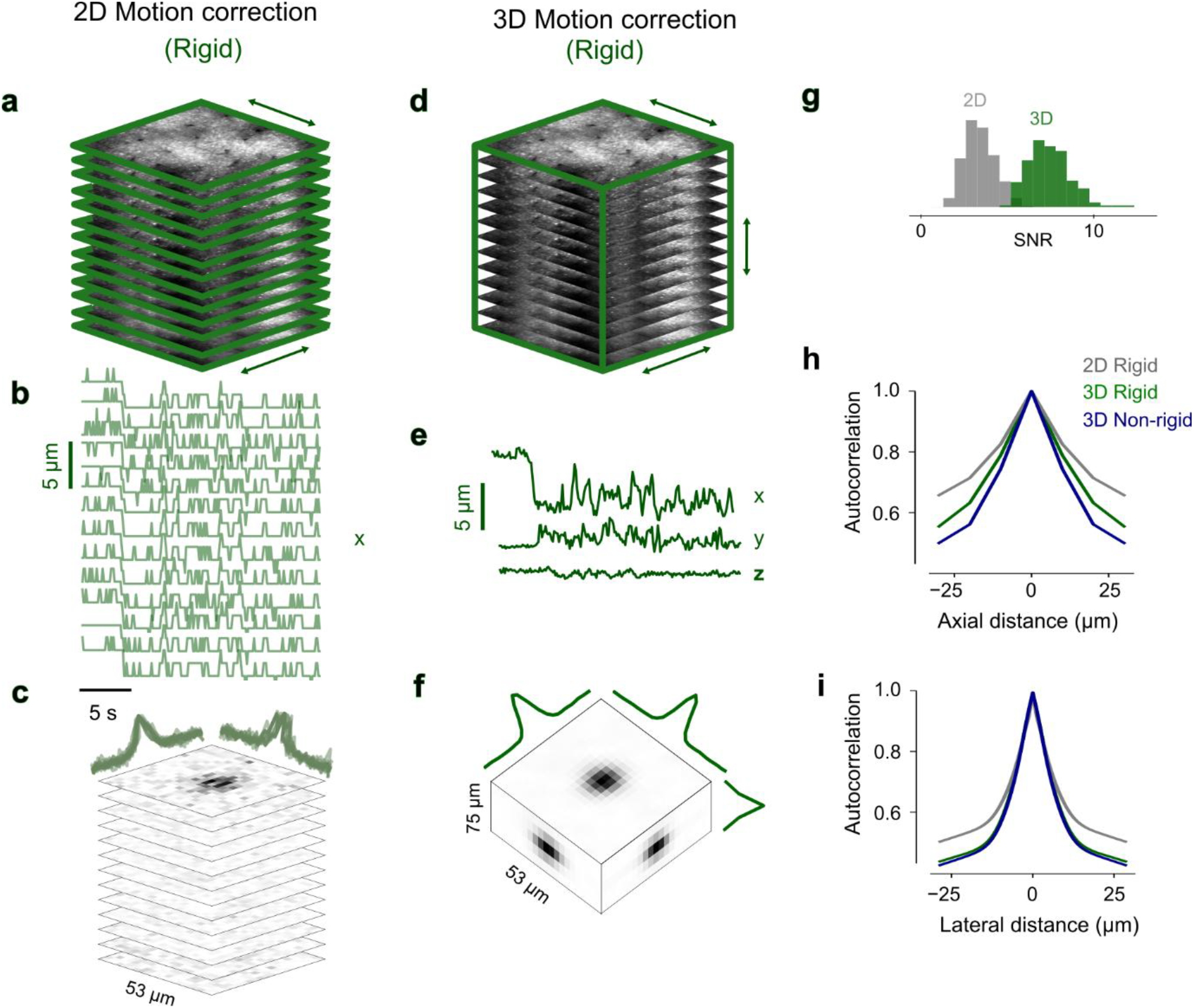
Comparison of 2D and 3D rigid motion correction approaches. **a**. Reference image for an example LBM recording. 2D rigid correction shifts plane independently, only correcting within-plane motion. **b**, Rigid motion estimates in the x direction for the stack of frames shown in (a) estimated with 2D motion correction. **c**, Phase correlation for an example frame for the planes in (a). Overlaid lines show the center slice for each patch. **d**. 3D rigid motion correction estimates three-dimensional motion for all planes simultaneously. **e**, Estimated x, y and z shifts for the volume shown in (d). **f**, 3D phase correlation map from a single frame for the example volume in (d). Curves indicate center slices along x, y, and z. **g**, For a single recording, distribution of the phase correlation SNR across planes (2D) and volumes (3D). **h**. Spatial autocorrelation in z of the motion-corrected movie. **i**. Same as (k), in x/y.

**ED Figure 2.**
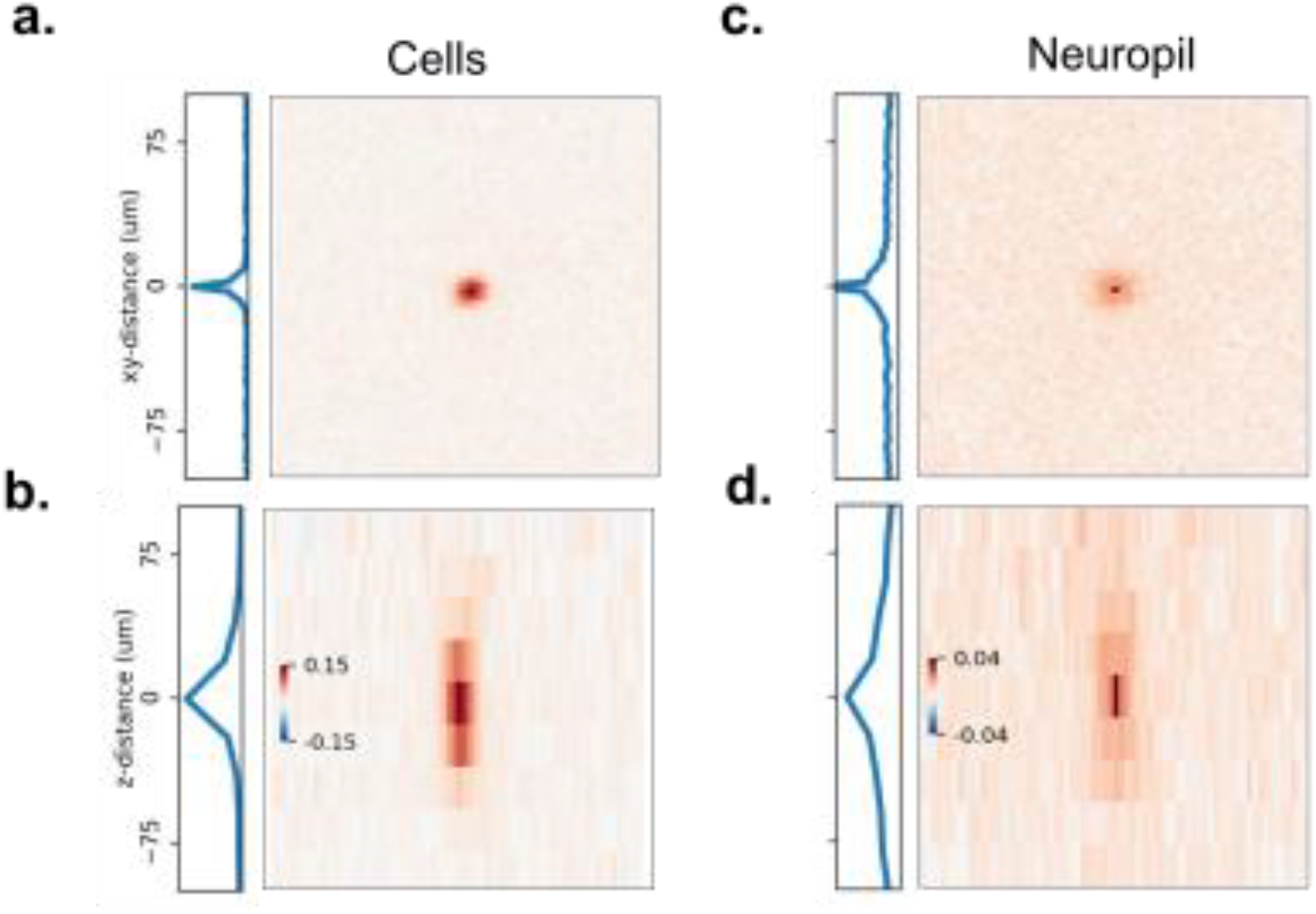
Volumetric data contains 3D correlation structures. **a**. The average fluorescence correlation of a voxel in a cell with its neighbors in x and y for an LBM recording. **b**. Same as (a) in x and z directions. **c**. Average fluorescence correlation of a neuropil voxel (e.g. a voxel not located within a cell) with its neighbors in x and y for the same recording as (a). **d**. Same as (c), in x and z.

**ED Figure 3.**
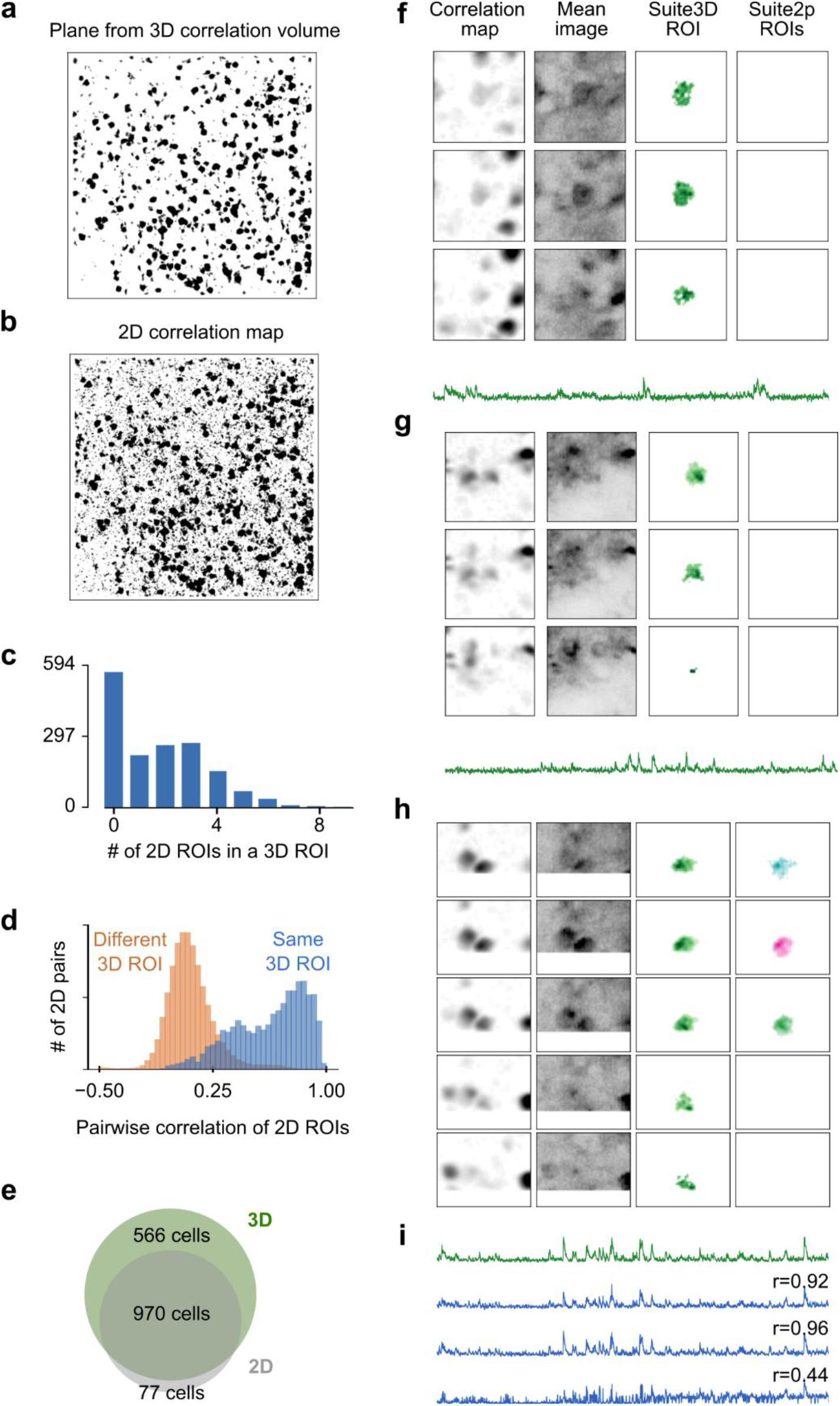
Comparison of detection and segmentation in 2D and 3D. **a**. An example plane of a 3D correlation volume computed from a 30 s recording in visual cortex (500 μm width). Linear colormap from 10^th^ to 90^th^ percentile. **b**. 2D correlation map for the same plane as (a). **c**. Histogram of the number of 2D ROIs per 3D ROI in the example recording. **d**. Histogram of pairwise correlation of 2D ROIs belonging to the same 3D ROI (blue) and different 3D ROIs (orange) overlap with no hard threshold. **e**. 3D detection finds almost all 2D ROIs, as well as many ROIs not found in 2D detection. **f,g**. Example cells across three planes detected in 3D detection and not in 2D. **h**. Example cell detected across five planes in 3D, but only on three planes in 2D. **i**. Trace of the 3D ROI (*green*) and the detected 2D ROIs (*blue*), each labelled with their correlation coefficient with the 3D trace.

**ED Figure 4.**
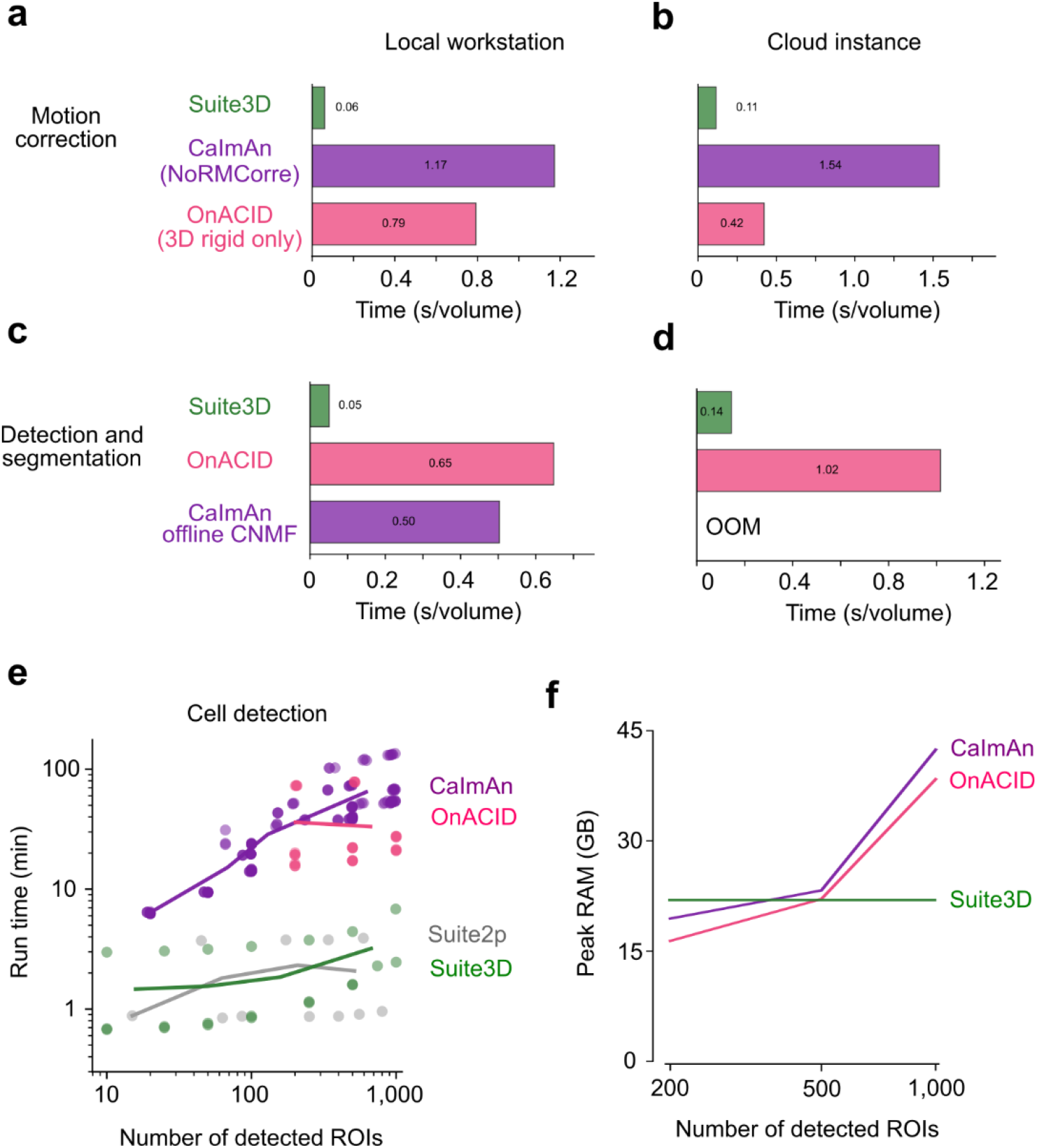
Runtime and memory requirements of tested algorithms. All benchmarking was conducted on a 6 x 512 x 512 dataset with 2,220 frames in V1. **A**. Time required to motion-correct a single acquired volume on a local workstation using three different pipelines. **b**. Same, on a cloud instance. **c**. Time required to go from a motion-corrected movie to segmented ROIs on a local workstation. **d**. Same, on a cloud instance. OOM stands for “out of memory”. **e**. Detection and segmentation runtime as a function of the number of ROIs detected on three datasets. Each point represents a unique parameter configuration and dataset, and the lines correspond to local means. Experiment conducted on a local workstation **f**. Peak RAM usage as a function of detected ROIs for one dataset, computed on a cloud instance.

**ED Figure 5.**
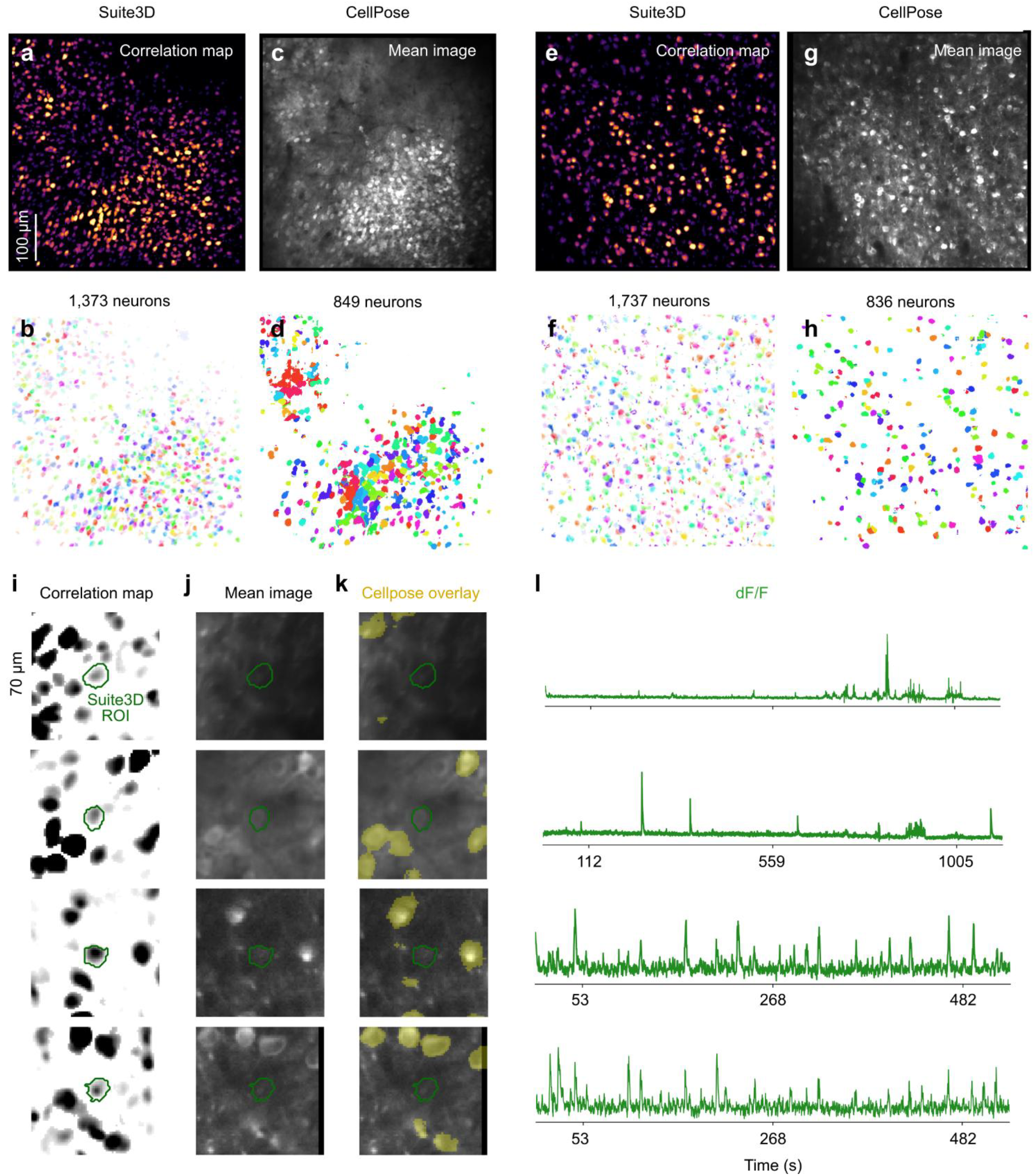
Comparison of Suite3D with CellPose. CellPose misses sparsely active cells, and struggles with densely packed FOVs. **a**. Example Suite3D correlation map from a CA1 recording. **b**. 1,373 neurons are detected in the full volume, and a single plane is shown here. **c**. Mean image from the same recording is fed into CellPose. **d**. CellPose ROI masks for a single plane, out of 849 total detected. **e-h** Same as a-d, for a field of view in mouse visual cortex. **i**. Four example Suite3D ROIs from CA1 (top 2) and V1 (bottom 2), overlaid on a 3D correlation map. **j**. Mean image for the same ROIs in (i). **k**. Overlay of CellPose ROIs detected from the mean images in (j). **l**. Suite3D-extracted traces of the example cells, which show sparse activity.

**ED Figure 6.**
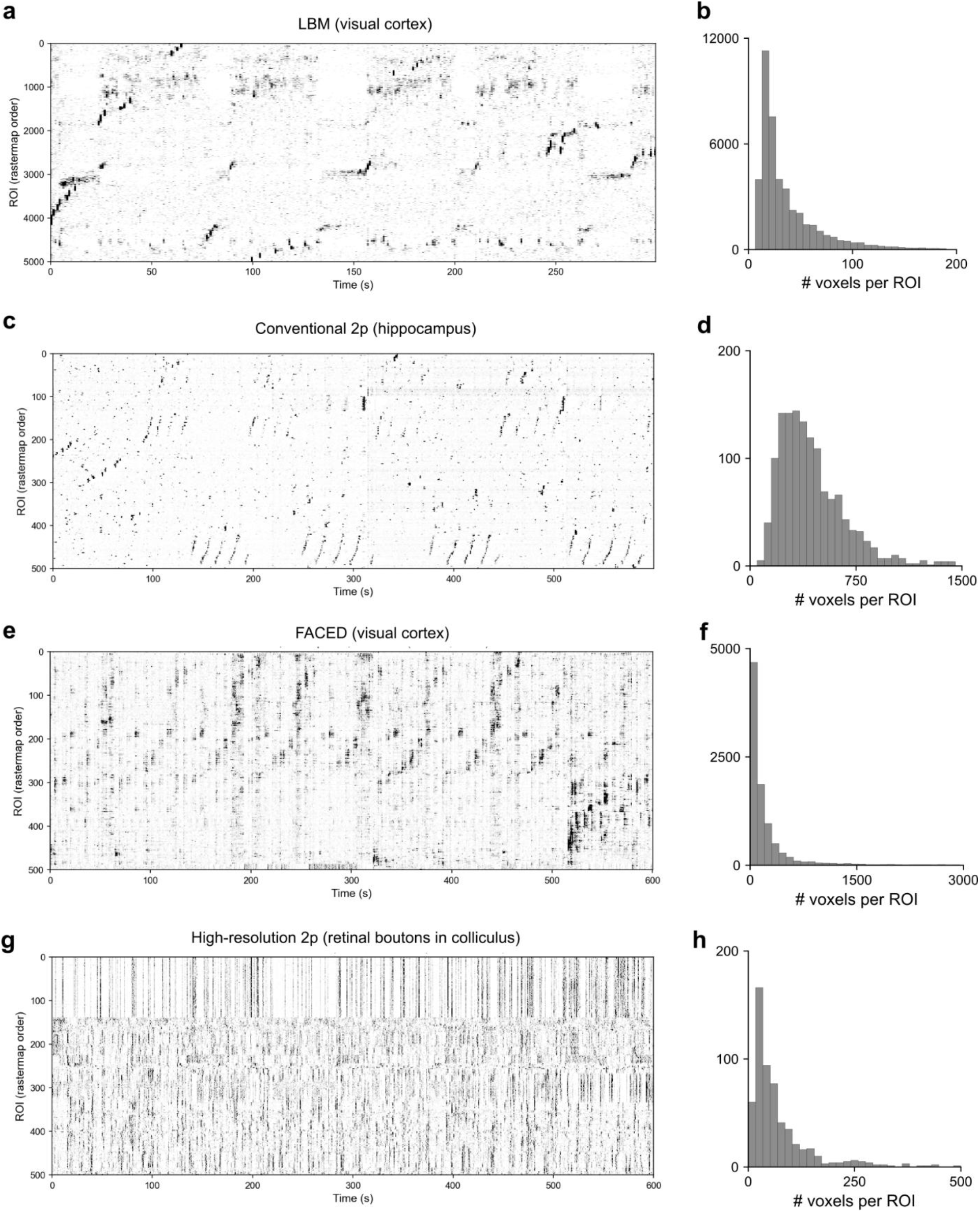
Validation on multiple datasets from Figure 5. **a**. Raster of deconvolved population activity for a subset of 500 cells identified by Suite3D in an LBM recording of visual cortex during spontaneous activity. Sorting the neurons with RasterMap^57^ reveals coordinated patterns of population activity. **b**. Number of voxels in each detected ROI. **c,d**. Same, for hippocampus neurons recorded during navigation in virtual reality. The neurons activated in sequences likely correspond to place cells. **e,f**. Same, for visual cortex neurons recorded with FACED during visual stimulation with gratings. The raster reflects the trial structure and differing selectivity of neurons. **g,h**. Same, for axonal boutons measured in the superior colliculus. The raster shows a mix of synchronized and desynchronized activity.

**ED Figure 7.**
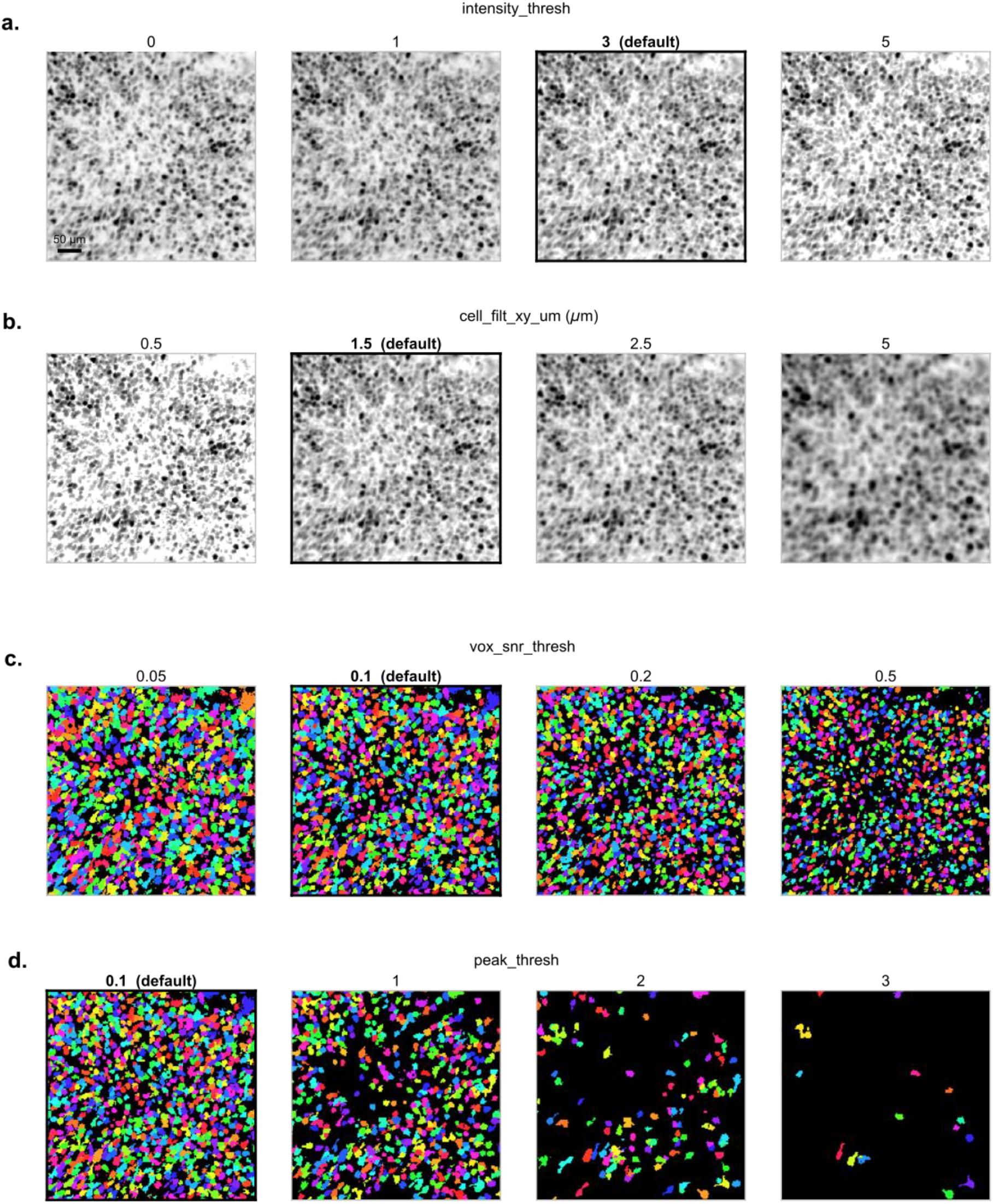
Parameter sweeping on a single dataset. On the hippocampus dataset, we used the sweep interface of Suite3D to vary one parameter at a time (see ED Table 2 for parameter descriptions). **a**. Lowering the intensity threshold increases the background level of the correlation volume – this parameter has a stronger effect on noisier recordings. **b**. Increasing the cell filter size reduces the amount of detail in the correlation volume. **c**. Increasing the voxel SNR threshold reduces the size of ROIs extracted. **d**. Increasing the peak threshold reduces the number of ROIs extracted.

**ED Figure 8.**
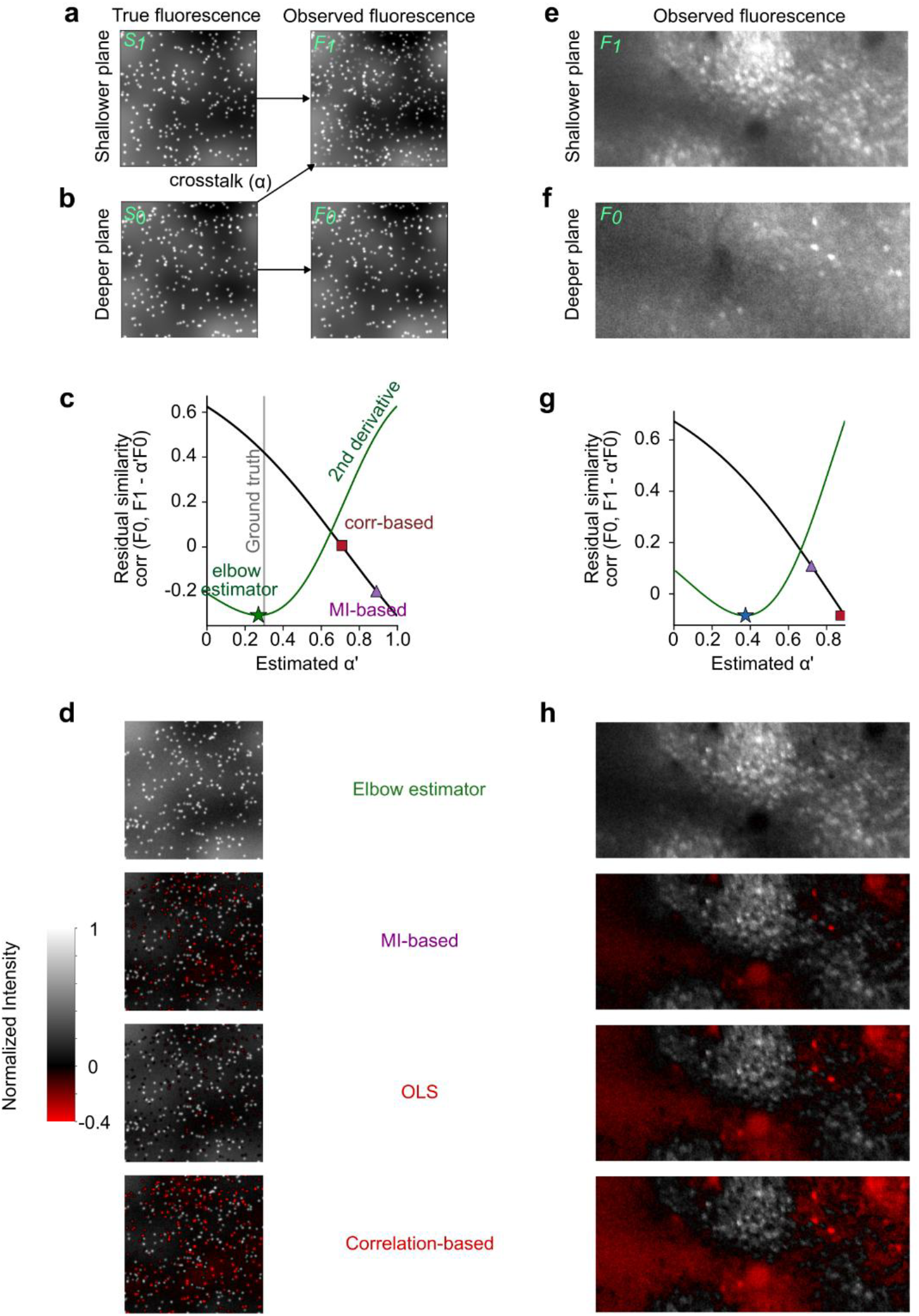
Method to subtract crosstalk in LBM recordings. **a,b**. Simulated data with low spatial frequency anatomical features and cells. The observed fluorescence in shallow planes (a) is contaminated by the crosstalk from deeper planes with weight α. Observed fluorescence in deeper planes is unaffected. **c**. The similarity of crosstalk-corrected fluorescence of the shallow plane with the deeper plane decreases as the subtracted component increases. Estimators that minimize correlation or mutual information (MI) between the shallow and deep planes overestimate the true crosstalk coefficient. **d**. Crosstalk-corrected fluorescence using the 2^nd^ derivative (elbow) estimator, MI- and correlation-minimizing estimators, and an ordinary linear regression of the shallow plane and deep plane. Non-elbow estimators over-subtract, and remove the true anatomical signal from the shallow plane, and create negative spots (red). **e,f**. Observed fluorescence from a LBM recording. **g,h** same as (c,d) on LBM data.

**ED Figure 9.**
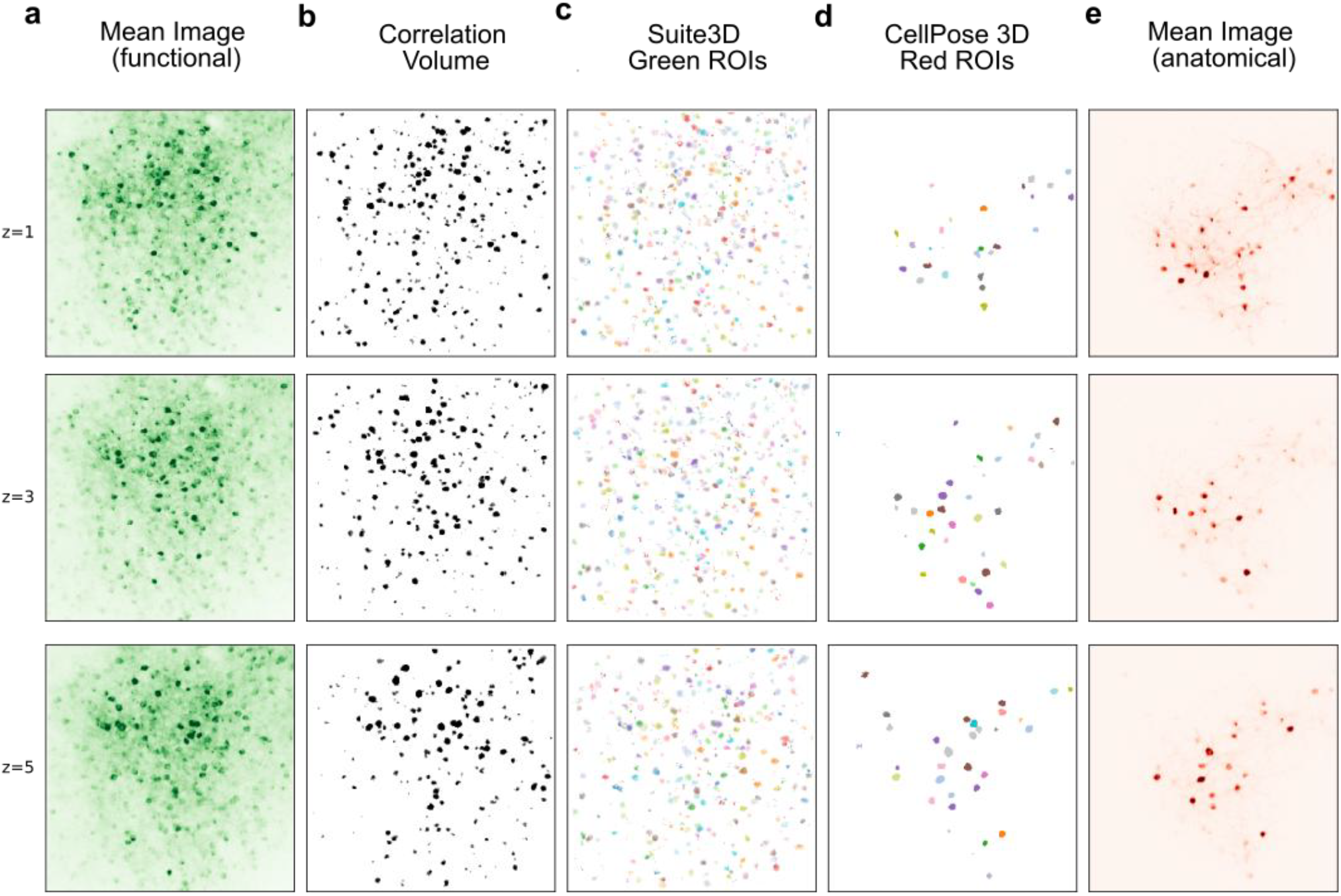
Ground-truth validation of cell masks with sparse anatomical labeling. **a**. Mean image of the motion-corrected green channel of a visual cortex recording. Rows correspond to 3 example planes, spaced 20 um apart. **b**. Correlation map computed with Suite3D. **c**. ROIs computed with Suite3D. **d**. Ground truth anatomical ROIs extracted with CellPose, which are used to evaluate the Suite3D ROIs. **e**. Motion-corrected mean image of the red anatomical channel used to extract the ground truth ROI masks.

**ED Table 1.** More details on the datasets, same order as Table 1.

| Fig. | Vox. Spacing (μm) | Vol. Rate (Hz) | # voxs (z,y,x, [total]) | # vols | # ROIs | Mean ROI size (z,y,x; μm) | Non-de-fault params | MC time (min) | Det. + seg. time (min) |
| --- | --- | --- | --- | --- | --- | --- | --- | --- | --- |
| 2dih, 3,4 | 10 x 1 x 1 | 3.34 | 7 x 515 x 514 (1,852,970) | 2,001 | 864 | 31 x 14 x 15 | - | 2.2 | 1.0 |
| 2a-c, e-g, i, k, l; 5a-c | 20 x 3.3 x 3.3 | 4.12 | 22 x 928 x 735 (15,005,760) | 1,235 | 43,652 | 65 x 20 x 20 | 3* | 5.1 | 11.8 |
| 5d-f | 10 x 1 x 1 | 6.00 | 4 x 514 x 512 (1,052,672) | 3,603 | 1,371 | 38 x 20 x 19 | - | 1.8 | 1.1 |
| 5g-i | 5 x 2 x 1.4 | 4.19 | 40 x 512 x 803 (16,445,440) | 2,516 | 9,323 | 33 x 20 x 16 | - | 28 | 16.2 |
| 5j-l | 2 x 1 x 1 | 7.27 | 6 x 256 x 259 (397,724) | 4,364 | 587 | 7 x 7 x 8 | 4** | 0.8 | 0.4 |

\*: intensity\_thresh=1 and vox\_snr\_thresh=0.05 to lower signal thresholds as LBM recordings have lower signal-to-noise ratio; ext\_subtract\_iters=0 due to large voxel size.
\*\* cell\_filt\_z\_um=1, cell\_filt\_xy\_um=1, npil\_filt\_xy\_um=70, npil\_filt\_z\_um=0; all to account for smaller features.

**ED Table 2.** Tunable parameters of Suite3D.

|  | Parameter | Default | Typical Range | Description |
| --- | --- | --- | --- | --- |
| **Motion Correction** | 3d_reg | True | True / False | Estimate rigid and non-rigid motion in three dimensions |
|  | nonrigid | False | True / False | Correction of non-rigid motion by 2D patches or 3D blocks (if 3d_reg=True) |
|  | max_rigid_shift_pix | 100 | 1–150 | Maximum number of pixels to consider for possible rigid shifts in x/y |
|  | block_size_3d | (5, 128, 128) | z 2–5, xy 32–256 | Size of the nonrigid blocks (z, y, x). |
|  | apply_z_shift | True | True / False | Apply the measured rigid z drift. For shallow stacks, or for recordings where z-estimation is poor, set to False |
| **Detection** | intensity_thresh | 5 | 0–5 | Minimum intensity of a single timepoint of a voxel on the detection movie required to contribute to the correlation volume. Reducing it increases the background of the correlation map, increasing it sparsifies the correlation map |
|  | detection_timebin | Auto | 1–10 | Temporal binning applied to the detection movie. Higher improves segmentation for noisier movies, at risk of missing cells with very rare and fast transients. |
|  | cell_filt_xy_um | 1.5 | 1–5 | Width (μm) of the cell filter. Setting it larger will exclude smaller neurites. |
|  | cell_filt_type | ‘gaussian’ | ‘gaussian’, ‘unif’ | Uniform or gaussian filter is applied to the correlation map. If applying uniform, typically need to increase extent by 2x. |
|  | npil_filt_xy_um | 100.0 | 50–100 | Width (μm) of the neuropil (background) filter subtracted before detection. |
|  | sdnorm_exp | 0.9 | 0.5–1.0 | Normalization exponent for the correlation map. For noisy recordings, reduce this to remove bright patches in dark areas. |
| **Segmentation** | peak_thresh | 0.1 | 0.03–0.15 | Minimum value of the correlation map to seed a new ROI. Reducing will increase number of ROIs detected, potentially including junk. Can also be filtered post-hoc from stats.npy |
|  | vox_snr_thresh | 0.05 | 0.05–0.20 | Fraction of a voxel’s variance the ROI’s trace must explain for the voxel to join the ROI. Reducing it creates larger ROIs, increasing creates smaller ROIs. |
|  | segmentation_spatial_filt | 2 | 0–5 | xy uniform smoothing (pixels) applied to the detection movie for segmentation. For noisy recordings, increasing creates smoother ROI masks. |

